# EUGENe: A Python toolkit for predictive analyses of regulatory sequences

**DOI:** 10.1101/2022.10.24.513593

**Authors:** Adam Klie, Hayden Stites, Tobias Jores, Joe J Solvason, Emma K Farley, Hannah Carter

**Author notes:** Correspondence to Hannah Carter.

## Abstract

Deep learning (DL) has become a popular tool to study cis-regulatory element function. Yet efforts to design software for DL analyses in genomics that are Findable, Accessible, Interoperable and Reusable (FAIR) have fallen short of fully meeting these criteria. Here we present EUGENe (**E**lucidating the **U**tility of **G**enomic **E**lements with **Ne**ural Nets), a FAIR toolkit for the analysis of labeled sets of nucleotide sequences with DL. EUGENe consists of a set of modules that empower users to execute the key functionality of a DL workflow: 1) extracting, transforming and loading sequence data from many common file formats, 2) instantiating, initializing and training diverse model architectures, and 3) evaluating and interpreting model behavior. We designed EUGENe to be simple; users can develop workflows on new or existing datasets with two customizable Python objects, annotated sequence data (SeqData) and PyTorch models (BaseModel). The modularity and simplicity of EUGENe also make it highly extensible and we illustrate these principles through application of the toolkit to three predictive modeling tasks. First, we train and compare a set of built-in models along with a custom architecture for the accurate prediction of activities of plant promoters from STARR-seq data. Next, we apply EUGENe to an RNA binding prediction task and showcase how seminal model architectures can be retrained in EUGENe or imported from Kipoi. Finally, we train models to classify transcription factor binding by wrapping functionality from Janngu, which can efficiently extract sequences in BED file format from the human genome. We emphasize that the code used in each use case is simple, readable, and well documented (https://eugene-tools.readthedocs.io/en/latest/index.html). We believe that EUGENe represents a springboard toward a collaborative ecosystem for DL applications in genomics research. EUGENe is available for download on GitHub (https://github.com/cartercompbio/EUGENe) along with several introductory tutorials and for installation on PyPi (https://pypi.org/project/eugene-tools/).

## Introduction

Cracking the cis-regulatory code that governs gene expression remains one of the great challenges in genomics research. Since the completion of The Human Genome project^1^, we have witnessed efforts to annotate the human genome that have generated an immense amount of functional genomics data^2,3^ and candidate regulatory elements^4^. This data has in turn powered machine learning methods aimed at predicting the functional readouts of these sequences such as histone marks^5^, chromatin accessibility^6^, 3D conformation^7^, and gene expression^8^. Deep learning (DL) has become especially popular in this space, and has been successfully applied to tasks such as DNA and RNA protein binding motif detection^9–12^, chromatin state prediction^13–23^, transcriptional activity prediction^16,24–27^ and 3D contact prediction^28,29^. Recently, complementary models have been developed to predict data from massively parallel reporter assays (MPRAs) that directly test the gene regulatory potential of candidate elements^30–32^. Most encouragingly, many of these multilayered models go beyond state of the art (SOTA) predictive performance to generate expressive representations of the underlying sequence that can be interpreted to better understand the cis-regulatory code^18,23,32^.

Despite these advances, executing a deep learning workflow in genomics remains a considerable challenge. Though model training has been substantially simplified by the continued development of dedicated DL libraries such as PyTorch^33^ and Tensorflow^34^, training nuances specific to genomics data along with complex preprocessing and interpretation methods create an especially high learning curve for performing analyses in this space. Though these libraries have built-in support for methods and visualizations of image and text-based data, utilities to handle genomics data are lacking. On top of this, the heterogeneity in implementations of most code associated with publications greatly hinders extensibility and reproducibility. These conditions often make the development of genomics DL workflows painfully slow even for experienced DL researchers and potentially inaccessible to many others.

Accordingly, the genomics DL community has assembled several software packages^35–40^ that each aim to address one or more of these challenges. However, each toolkit on its own does not offer both comprehensive functionality and simplicity, and there remains a general lack of interoperability between packages that is essential for sustained improvement and utility in the fast-advancing field of DL. For instance, Kipoi^37^ greatly lowers the accessibility barrier to trained models and published architectures, but does not provide a comprehensive framework for an end-to-end DL workflow. Selene^36^ implements a library based in PyTorch for applying the full DL workflow to new or existing models, but offers a limited programmatic interface and requires the use of complex configuration files. Janggu^38^, one of the more comprehensive of the available packages, provides extensive functionality for data loading and for training Keras models, but offers limited support for PyTorch and limited functionality for model interpretation. More recently, ENNGene^39^ was designed to create a simple graphical user interface (GUI) for non-computational users, but offers limited programmatic customizability for more advanced users. Generally, there is a need for a *comprehensive* toolkit in this space that follows FAIR data and software principles^41,42^ and that is inherently designed to be *simple* and *extensible*.

Here we introduce EUGENe (**E**lucidating the **U**tility of **G**enomic **E**lements with **Ne**ural Nets), a FAIR toolkit for the analysis of sequence-based datasets modeled after Scanpy^43^. In this work, we first summarize the key functionality of the EUGENe package by describing each of the modules it contains and their guiding principles. We then describe the two fundamental data structures that give EUGENe its simplicity and extensibility in detail, namely SeqData and BaseModel. Finally, we show the *application* of EUGENe to three separate sequence prediction tasks: promoter activity prediction in plants, *in vitro* RNA binding prediction with SOTA model architectures, and transcription factor binding classification from ChIP-seq data. In each, we demonstrate the ability of simple and well-documented EUGENe code to achieve high predictive performance and biological interpretability.

## Results

### The EUGENe workflow

A standard EUGENe workflow consists of the 3 main stages outlined in **Figure 1**: extracting, transforming and loading (collectively ETL) data from common file formats (**Figure 1a)**, instantiating, initializing and training (collectively IIT) neural network architectures (**Figure 1b)**, and evaluating and interpreting (EI) learned model behavior on held-out data (**Figure 1c**). The EUGENe package implements this workflow through several modules. Each module implements functionality that is principally designed to be self-contained given the proper input data, but that can interface with one or more other modules. We designed the workflow to be run within a notebook interface via the EUGENe Python application programming interface (API). We discuss the purpose and functionality of each module briefly below. For more information, see the tool’s documentation pages (https://eugene-tools.readthedocs.io/en/latest/).

**Figure 1.**
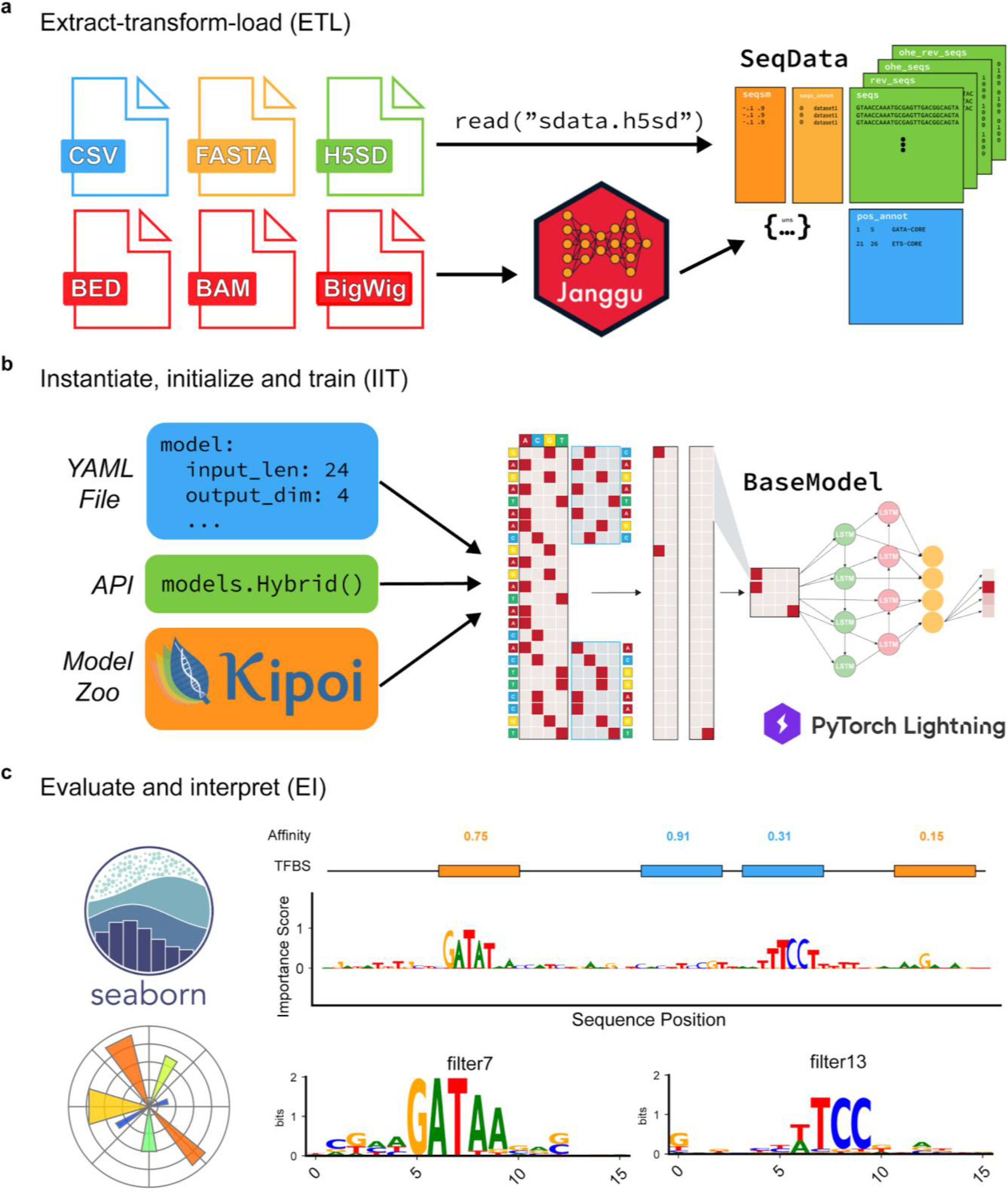
EUGENe workflow for predictive analyses of regulatory sequences. The EUGENe workflow can be broken up into three primary stages: **a**, data extraction, transformation and loading (ETL), **b**, model instantiating, initializing and training (IIT), and **c**, model evaluation and interpretation (EI).

EUGENe currently supports loading DNA or RNA sequence inputs from CSV, NumPy, FASTA, and a custom h5 file implementation we named H5SD (**Figure 1a**, *dataload* module). We also have wrapped functionality from the Janggu package^38^ for reading sequences from BED, BAM and BigWig files (*external* module, see **JunD ChIP-seq binding classification** section for details). On top of allowing users to load their own datasets, we supply a collection of hand-curated benchmarking datasets that are readily available for download and subsequent data loading with a single function call (*datasets* module, **Supplementary Table 1**). All sequences and metadata are loaded into a standardized SeqData format that EUGENe functions act on to perform many standard per sequence preprocessing tasks (*preprocess* module) like reverse complementation or one-hot encoding, as well as whole dataset functions like train and test set splitting (e.g. by chromosome or by fraction) and target variable normalization (e.g. z-score, clamping, etc.). EUGENe also offers functions for converting preprocessed data into training-ready formats (e.g. PyTorch dataloaders) from SeqData and other more general Python objects (e.g. NumPy arrays and Pandas DataFrames).

EUGENe provides three classes of model architecture (*models* module) for users to work with: Base, SOTA and Custom (**Supplementary Table 2)**. As Base Models, we currently offer built-in and customizable fully connected (FCN), convolutional (CNN), recurrent (RNN) and hybrid (a combination of the three, **Figure 1b**) architectures that can be specified to incorporate information from both the forward and reverse strand. We also offer simple functions for instantiating two customizable architectures often used in benchmarking tasks (SOTA Models), DeepBind and DeepSEA. Finally, we offer a route for users to design their own custom architectures (Custom Models) that requires the implementation of only two functions (instantiation and forward propagation) and provide a tutorial that walks through this process on EUGENe’s GitHub (https://github.com/cartercompbio/EUGENe/blob/main/tutorials/adding_a_model_tutorial.ipynb). To train instantiated models and handle standard tasks like optimizer configuration and metric logging, EUGENe relies on the PyTorch Lightning (PL) framework (*train* module). Though EUGENe’s PL wrappers offer valuable abstraction from many boilerplate implementation tasks, the framework also gives users the flexibility necessary to design custom architectures and training schemes (e.g. custom optimizers, loss functions, etc.) if desired. We also provide a set of wrapper functions for utilizing trained PyTorch and Keras models and architectures from the Kipoi model zoo^37^ (*external* module).

Interpretation of trained models has been crucial for deciphering aspects of the cis-regulatory code and is a core aspect of the EUGENe workflow (*interpret* module, **Figure 1c**). There are many strategies for model interpretation in genomics^44–51^, but three categories are repeatedly used and thus implemented in EUGENe: filter visualization, feature attribution and *in silico* experimentation (**Supplementary Figure 1**). Filter visualization is applicable to model architectures that begin with a set of convolutional filters and involves using the set of sequences that significantly activate a given filter (maximally activating subsequences) to generate a position frequency matrix (PFM) (**Supplementary Figure 1a**). Multiple methods exist for choosing the maximally activating subsequences and we have implemented two of them so far in EUGENe^9,17^. The PFM can then be converted to a position weight matrix (PWM), visualized as a sequence logo and annotated with tools like TomTom^52^ using databases of known motifs such as JASPAR^53^ or HOCOMOCO^54^. Feature attribution involves using the trained model to score every nucleotide of the input on how it influences the downstream prediction for that sequence **(Supplementary Figure 1b**). In EUGENe, we currently implement or integrate several common feature attribution approaches, including standard *in silico* saturation mutagenesis (ISM), InputXGradient^55^, DeepLIFT^55^ and GradientSHAP^56^. Finally, EUGENe offers a simple set of functions that use trained models as *in silico* oracles to perform sequence evolution and feature implantation experiments (**Supplementary Figure 1c)**.

Data visualization is another key component of the EUGENe workflow (*plotting* module). We provide a large suite of functions for performing exploratory data analysis, generating performance summaries and visualizing model interpretations that all utilize the Matplotlib ^57^ and Seaborn^58^ libraries. This gives users the flexibility to customize the plots they generate with EUGENe.

### Building workflows through SeqData and BaseModel

To promote a streamlined data analysis process, we introduce SeqData (**Figure 2a**), a Pythonic data structure modeled after the popular AnnData used in the single cell field^59^. Almost all functionality in EUGENe is designed to act on and modify these standardized objects that act as organized containers for DNA and RNA sequences (*seqs, rev_seqs*), sequence representations (*ohe_seqs, ohe_rev_seqs*), and sequence annotations (*seqs_annot*, e.g. training targets), Data are loaded into SeqData by pointing to files on disk or by calling for a specific dataset defined in the *datasets* module (**Figure 2a**). Once created, an array of functions can be called directly on these objects to perform preprocessing (**Figure 2a)**, data visualization (**Figure 2b**) and data conversion to formats ingestible by deep learning frameworks (**Figure 2c**). Trained models can be used to make predictions on sequences stored in SeqData, and these predictions are stored and automatically accessed by functions that generate performance visualizations. Filter visualization, feature attribution and *in silico* experimentation can all be run through SeqData objects, generating per sequence and per dataset features stored in the sequence metadata and unstructured data (*uns*) attributes. These features can then be used to generate powerful sequence visualizations such as sequence and motif logos (**Figure 2d**) or dimensionality reduced clusterings (the latter stored in the object’s multidimensional attribute *seqsm*). Furthermore, SeqData and functions that act on SeqData are intentionally implemented as simple wrappers around widely used Python data structures (NumPy arrays, Pandas DataFrames, etc.) to enable the user to utilize the functionality of many standard Python libraries.

**Figure 2.**
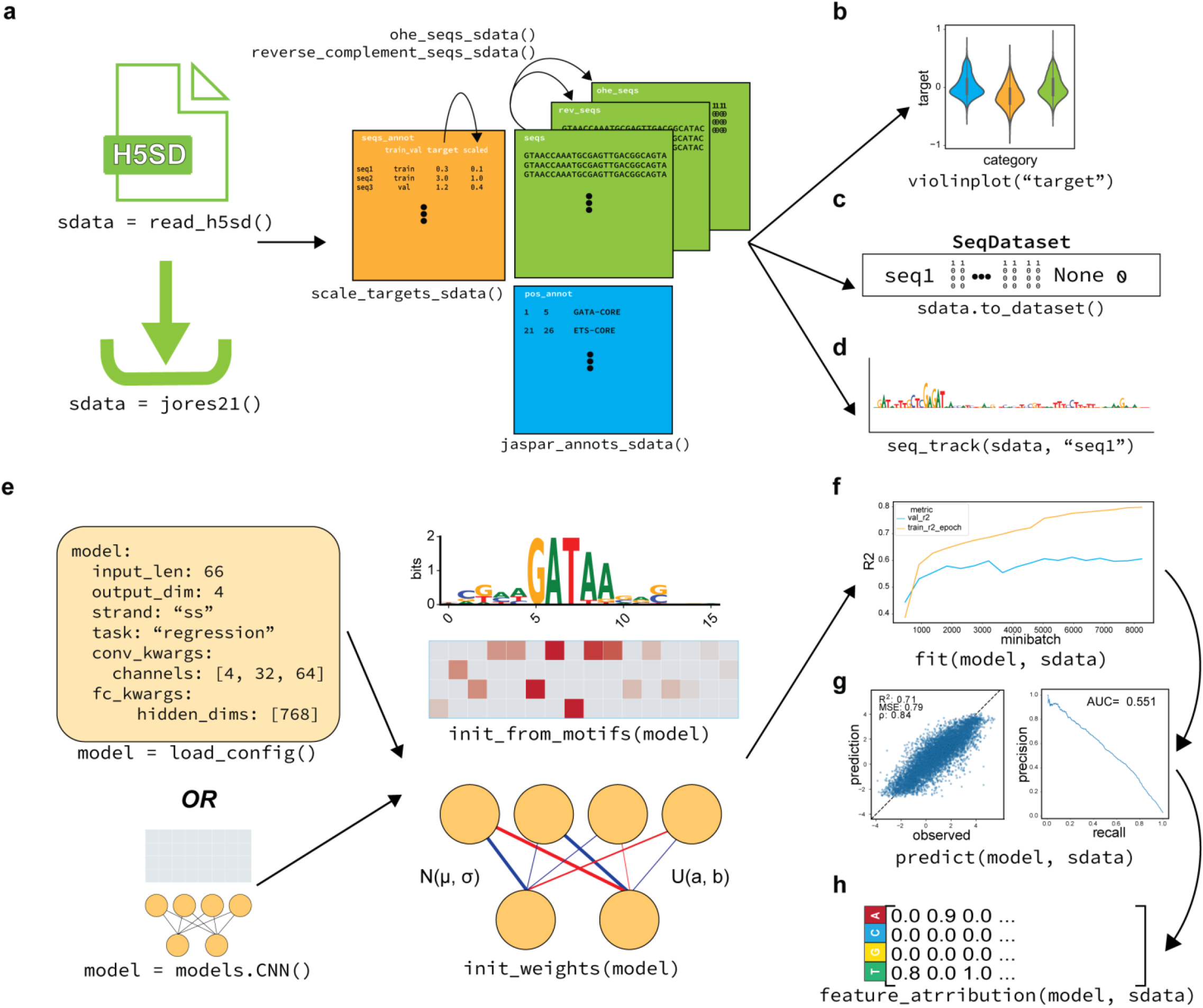
EUGENe maintains simplicity by operating on two fundamental objects: SeqData and BaseModel. **a**, SeqData objects can be read into memory from files already on disk, or by calling for a dataset available for download (`datasets.csv`). Once instantiated, SeqData objects containerize the EUGENe workflow, easing the preprocessing of sequences and of sequence metadata, **b**, the generation of exploratory data analysis plots, **c**, the creation of PL loadable datasets and objects, and **d**, the visualization of sequences and positional metadata as logos. **e**, A model can be instantiated either from a configuration file that specifies the hyperparameters of the model or from the API with hyperparameters passed in as arguments. Instantiated architectures can first be initialized with a desired initialization scheme, then **f**, fit to training data, **g**, used to predict on held-out data, and **h**, interpreted. Performance metric (right) training curves are pictured in **f**, test set performance curves for regression (left) and classification (right) are depicted in **g**, and a toy feature attribution matrix for a single sequence is depicted in **h**.

The standardized way of instantiating, initializing and training neural network architectures in EUGENe are BaseModel objects. We offer two main ways of instantiating model architectures: single function calls or configuration files with simple structure (**Figure 2e**, left). Custom architectures can also be imported from Kipoi or written from scratch with users only needing to define the architecture (*init* function) and the way forward propagation is handled (*forward* function) (**Supplementary Figure 2a**). After instantiation, models can be initialized with all the starting parameters sampled from popular distributions, or in the special case of convolutional filters, initialized with known motifs (**Figure 2e**, right). Once initialized, models can be fit to datasets (**Figure 2f**) by specifying the input sequence length, the number of outputs, the strand information to incorporate (i.e. whether to include reverse complement strand information and how to incorporate it) and the task type (e.g. regression versus classification). At a technical level, this is done using a PL implementation of BaseModel that handles the boilerplate aspects of model optimization, including but not limited to: optimizer and loss function configuration, training and validation set looping and metric logging. For many tasks, we find the built-in training scheme to be sufficient. However, due to the inherent flexibility of PL, more advanced users can customize almost all aspects of their model training strategy. For instance, custom training loops can be defined for models that implement multipart loss functions or use multiple optimizers^60–62^ (e.g. for variational autoencoders and generative adversarial networks respectively). (**Supplementary Figure 2b**). Once trained, models can be applied to held-out data to assess performance and generalizability (**Figure 2g)** or used as feature extractors for transfer learning approaches (**Supplementary Figure 2c**). Trained models can then be interpreted with function calls that store visualizable results directly in SeqData (**Figure 2d, h**).

### EUGENe use cases

#### STARR-seq plant promoter activity prediction

To showcase the functionality of EUGENe, we applied the toolkit to three example use cases, each of which highlights core aspects of the workflow on different data types and training tasks. We first used EUGENe to analyze published data from a STARR-seq assay of plant promoters^31^ (**Figure 3a**). In this work, Jores *et al* selected promoter sequences from −165 to +5 relative to the annotated TSS for protein-coding and mircoRNA (miRNA) genes of *Arabidopsis thaliana*, *Zea mays* (maize) and *Sorghum bicolor*. A total of 79,838 170-bp promoters were used to transiently transform two separate plant systems, tobacco leaves and maize protoplasts, and regulatory activity was quantified using plant STARR-seq^63^ in each system. These assays provide two activity scores that can serve as single task regression targets for training EUGENe models.

**Figure 3.**
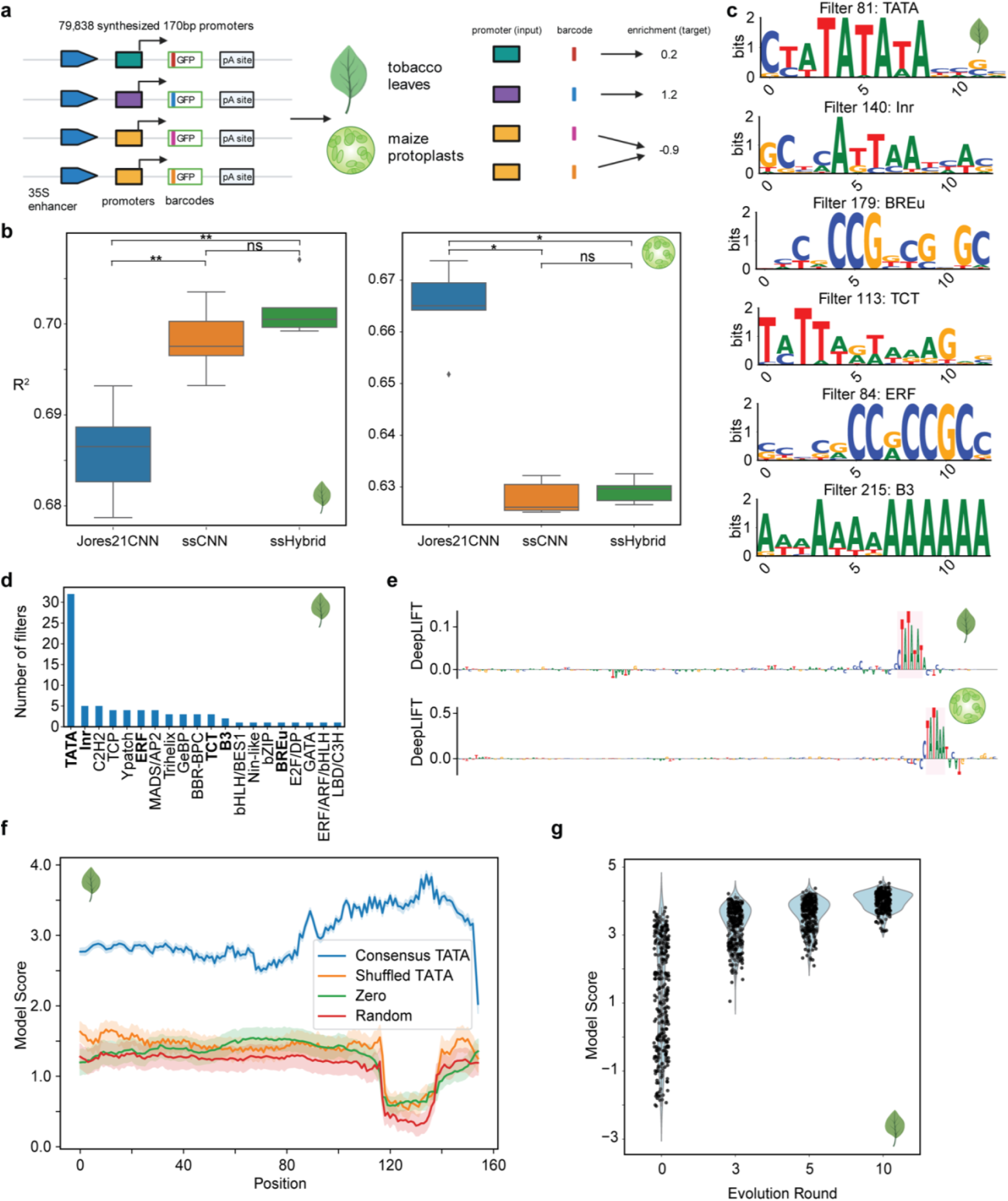
EUGENe models identify the TATA box and several TF motifs to accurately predict regulatory activity. **a**, jores21 use case schematic. We trained EUGENe models to predict the regulatory activity of 79,838 plant promoters quantified by plant STARR-seq in tobacco and maize. **b**, Performance comparison of three convolution-based architectures on predicting promoter activity in tobacco leaves (left) and maize protoplasts (right). The boxplots show distributions of R^2^ values on held-out test data for each architecture across 5 random initializations. **c**, A hand selected set of convolutional filters visualized as PWM logos that had significant annotations to known core promoter elements (CPE) and transcription factor (TF) binding clusters in plants **d**, Histogram showing the number of learned filters assigned to CPEs and TF binding clusters by TomTom with bolded annotations corresponding to the logos in **c**. **e**, Sequence logo visualizations of feature importance scores calculated using the DeepLIFT algorithm on the highest predicted test set sequence in the leaf (top) and protoplast (bottom) model. **f**, Model scores for 310 sequences implanted with a 16bp sequence containing a consensus TATA box motif, a shuffled version of the same sequence, an all zeros sequence and a random sequence (all 16bp in length). The 95% confidence interval is shown. **g**, Model scores for the same set of 310 promoters at different rounds of evolution compared against baseline predictions (evolution round 0). The best leaf model was used to generate panels **c, d, f** and **g** (protoplast model results are shown in **Supplementary Figure 3**). *p < 0.05, **p<0.01, ns = not significant. Mann-Whitney U test p-values corrected by the Benjamini-Hochberg method.

We first made this dataset readily loadable through the EUGENe *datasets* module (as `jores21`) and implemented both the custom BiConv1D layer^64^ and CNN architecture (Jores21CNN) described in Jores *et al*. We then trained separate Jores21CNN architectures for predicting tobacco leaf activity scores (leaf models) and maize protoplast activity scores (protoplast models) and benchmarked them against built-in CNN and Hybrid architectures with matched hyperparameters (**Supplementary Table 3**). To perform training as described in Jores *et al* (see **Methods**), we initialized 78 filters of the first convolutional layer of all models with position weight matrices of plant transcription factor (n=72) and core promoter element (n=6) PWMs^31^. The rest of the parameters of each model were randomly initialized 5 separate times and trained to assess reproducibility. In the leaf system, we noted similar performances across architectures on held-out test data (**Figure 3b)**, with the Hybrids and CNNs outperforming Jores21CNNs when evaluated by variance explained (R^2^). We observed the opposite trend for protoplast models, where Jores21CNNs performed better than built-in CNNs and Hybrids (**Figure 3b)**. In both systems, all performance metrics for the most predictive models were comparable to those reported in Jores *et al* (**Supplementary Figure 3a**, **Supplementary Table 4**). We also trained models on activity scores from both leaves and protoplasts (combined models) and noted a marked drop in performance (**Supplementary Figure 3b**), underscoring the differences in the way the leaf and maize systems interact with the same set of promoters^31^.

We next applied several of EUGENe’s interpretation functions to the trained models to determine the sequence features each used to predict plant promoter activity. First, we used a filter visualization approach^17^ to generate PWM representations for each of the first convolutional layer’s filters (**Supplementary Figure 1a**) and applied the TomTom tool to annotate them (**Supplementary Table 5**). We queried the PWMs against the 78 motifs used to initialize the convolutional layers, both to determine if the initialized filters retained their motifs and to see if randomly initialized filters learned them *de novo*. For the leaf model, many of the learned filters were annotated to the core promoter elements, including the TATA box binding motif that was assigned to 32 filters (**Figure 3cd**). No learned filters from the protoplast model were assigned a significant annotation by TomTom (**Supplementary Figure 3c**), consistent with the observed performance drop in this system (**Supplementary Figure 3a)**. Next, we applied the DeepLIFT method^65^ to determine the individual nucleotide contributions for each test set sequence prediction (**Supplementary Figure 1b**). For many of the sequences with the highest observed activity scores, the TATA box motifs were often the lone salient feature identified (**Figure 3e, Supplementary Figure 3d**). In fact, when only a TATA box motif was inserted into every possible position in each of 310 selected promoters (**Supplementary Figure 1c**), we observed an 142% average increase in predicted activity across insertion positions and sequence contexts for the leaf model (**Figure 3f, Supplementary Figure 3e**). We also noted that the magnitude of the increase was dependent on position of insertion^66^, with the highest increases in predictions observed directly upstream of the TSS (**Figure 3f, Supplementary Figure 3e**). Finally, we performed 10 rounds of *in silico* evolution on the same set of 310 promoters as described in Jores *et al* (**Supplementary Figure 1c**). Almost all starting promoters showed a significant increase in predicted activity after just three mutations (**Figure 3g, Supplementary Figure 3f**). These results showcase a representative example of the way EUGENe’s interpretation suite can be used to identify key features of the cis-regulatory code underlying gene expression.

#### *In vitro* RNA binding prediction with DeepBind

To highlight the versatility of EUGENe to handle different inputs and prediction tasks, we next applied the toolkit to analyze RNA binding protein (RBP) specificity data first introduced in Ray *et al*^67^ and analyzed with DL in Alipanahi *et al*^9^. In the latter work, they trained 244 CNN models (DeepBind models) that each predicted the binding patterns of a single RBP on a set of 241,357 RNA probes (**Figure 4a**). The full probe set was designed to capture all possible RNA 9-mers at least 16 times and was split into two balanced subsets, Set A and Set B, for training and validation respectively (see **Methods**)^67^. Each RBP was incubated with a molecular excess of probes from each subset (in separate experiments) and subsequently recovered by affinity purification. The RNAs associated with each RBP were then quantified by microarray and subsequent bioinformatic analysis^68^. This yielded a vector of continuous binding intensity values for each RBP across the probe set that can be used for prediction.

**Figure 4.**
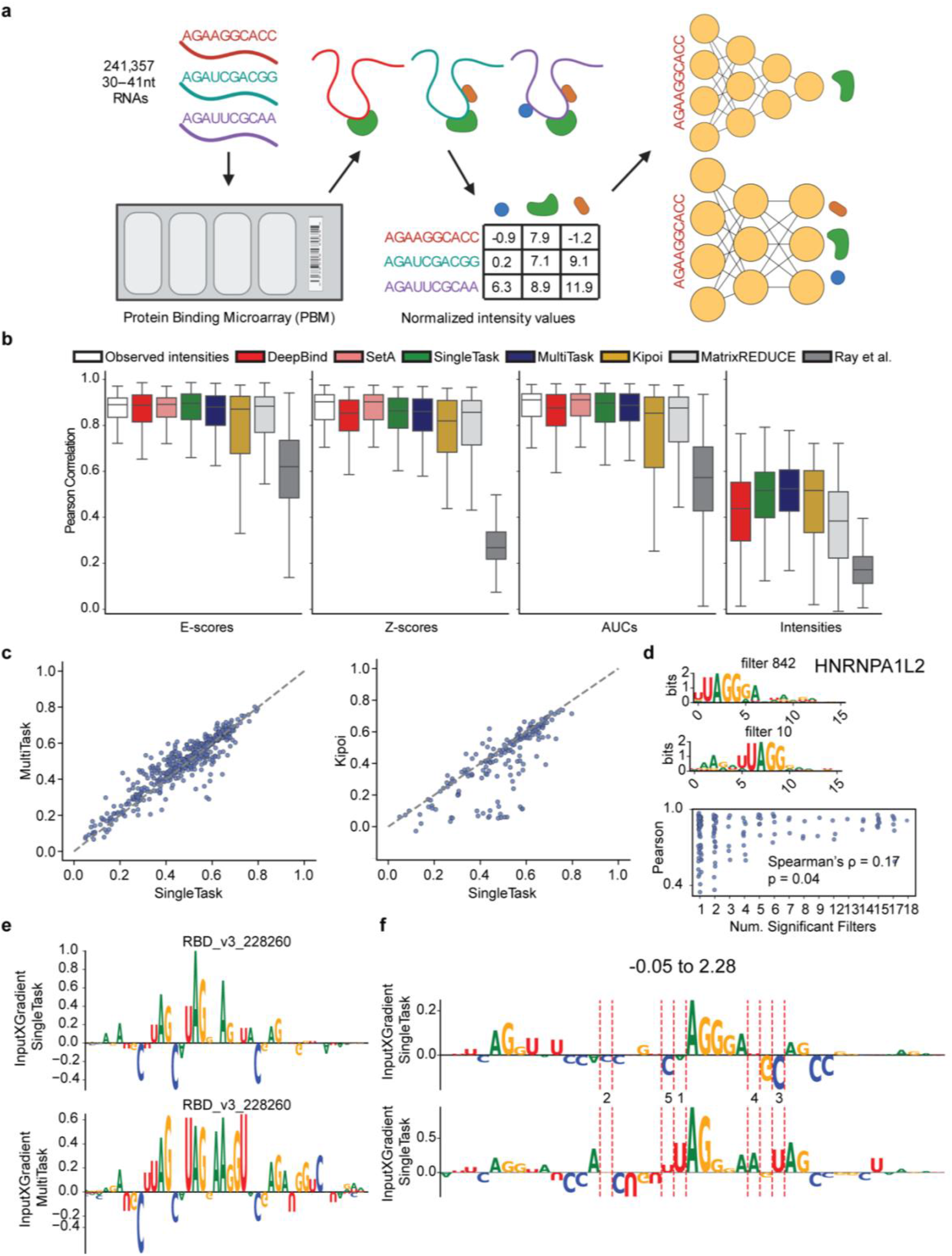
Prediction of RNA binding specificity with single task (ST), multitask (MT) and pretrained models (Kipoi). **a**, ray13 use case schematic.A set of 241,357 RNA probes were assayed against 244 RNA binding proteins (RBPs) to generate a 241,357 × 244 dimensional matrix of normalized intensity values. **b**, Pearson correlations across four different metrics with each metric calculated from comparisons between observed (Set B) and predicted binding intensities (see **Methods** for more details on how each metric is calculated). Each boxplot indicates a distribution of Pearson correlations across all 244 RBPs. Ray *et al*, MatrixREDUCE, DeepBind and Observed intensities refer to correlations calculated from predicted intensities reported in Alipanahi *et al*. Observed intensities and SetA refer to correlations calculated using the intensities from Set A probes as the predicted intensities (see **Methods**). **c**, Performance comparison scatterplots for ST models against MT models (left)and against Kipoi models (right). Each dot indicates a comparison of the Pearson correlation between predicted and observed intensities for two models on a single RBP. **d**, (top) A multitask filter with a TomTom significant annotation for HNRNPA1L2 visualized as a PWM logo. (middle) A filter for the single task HNRNPA1L2 model with a significant TomTom annotation for HNRNPA1L2. (bottom) The relationship between multitask performance (using the Z-scored Pearson correlations of observed and predicted intensities) on the y-axis, against the number of filters that were annotated with the corresponding RBP for that task on the x-axis. The Spearman’s correlation coefficient and associated p-value are shown. **e**, Feature attributions for the sequence with the highest observed intensity in the test set for HNRNPA1L2. The attributions were calculated using InputXGradient for single task (top) and multitask (bottom) models. **f**, The InputXGradient attribution scores for a random (top) and evolved (bottom) sequence after evolution with the HNRNPA1L2 single task model. Red dashed lines indicate mutations made during evolution and are annotated with the round the mutation occurred in.

To prepare for training, we first added this dataset to the *datasets* module (as `ray13`) and implemented a flexible DeepBind architecture in EUGENe (see **Methods**). We randomly initialized^69^ and trained 244 single task models using a nearly identical training procedure to Alipanahi *et al*. However, we used the Adam optimizer^70^ instead of the stochastic gradient descent algorithm and we used 32 filters in the convolutional layer instead of 16 (**Supplementary Table 6**). Along with these single task models, we also randomly initialized and trained a multitask model to predict 233 RBP specificities (i.e. a 233 dimensional vector) in a single forward pass, excluding 11 RBPs due to a high proportion of missing values across probes in the training set. Our multitask model had the same general architecture as the single task models (**Supplementary Table 6**), but with an increased number of convolutional filters (1024 as opposed to 32 for single task models) and a larger hidden layer size in the fully connected part of the model (512 as opposed to 32 for single task models). We also loaded 89 existing Kipoi^37^ models trained on a subset of human RBPs in the `ray13` dataset.

To evaluate model performance, we implemented functionality for calculating k-mer based Z-scores, AUCs and E-scores^9,67^ and added them to EUGENe’s metrics library in the *evaluate* module (see **Methods**). All models were trained on probe intensity measurements from Set A and evaluated with these metrics on measurements from Set B. By using k-mer based metrics, we can evaluate the concordance between Set A and Set B even though they include different sequence probes. We noted that the performance on Set B for all deep learning models was on par with Set B’s correlation to Set A (**Figure 4b, Supplementary Figure 4a**) and both single task and multitask models trained with EUGENe showed comparable performance to Kipoi and DeepBind models (**Figure 4bc**, **Supplementary Figure 4ab, Supplementary Table 7**). The reason for the poor observed performance of certain Kipoi models is not immediately clear, but could relate to differences in sequence or target preprocessing prior to evaluation. Though the ability to load these pretrained models from Kipoi is very useful for benchmarking, implementing and retraining models is usually necessary for fair comparisons of performance. EUGENe supports both loading and retraining models, allowing users to more quickly design and execute quality benchmarking experiments.

We also observed similar levels of performance of single and multitask models across metrics (**Figure 4bc, Supplementary Figure 4ab)**, consistent with the successful application of multitask models to the prediction of chromatin state from DNA input in bulk and single cells^13–23^. Using a multitask model offers much less training overhead and faster inference times than training 100s of single task models across tasks. Overall, in both the multitask and single task frameworks, we can train several high performing predictors of RBP specificity across more than 200 RBPs.

We next applied EUGENe’s interpretation suite to our trained models, first using the filter visualization approach outlined in Alipanahi *et al* to generate PFMs for convolutional filters. We again used TomTom to identify filters annotated with canonical RBP motifs^67^ in both the best performing single task models and the multitask model (**Figure 4d, Supplementary Figure 4c, Supplementary Table 8**) and found that the number of multitask filters annotated to an RBP was correlated with predictive performance for that RBP (**Figure 4d**). We also calculated feature attributions for all Set B sequences using the InputXGradient method and observed that canonical motifs were learned by both single task and multitask models (**Figure 4e, Supplementary Figure 4d**). Finally, we used EUGENe’s *in silico* functionality to evolve 10 random sequences using the single task HNRNPA1L2 model and visualized the feature attributions for these 10 sequences before and after five rounds of evolution. Several of the mutations that most increased the predicted score were those that generated canonical binding motifs for the protein (**Figure 4f**). We repeated this for two other RBPs (Pcbp2 and NCU02404) and observed that each model prioritizes mutations that create canonical binding motifs specific to the RBP they were trained on (**Supplementary Figure 4e**). Altogether, these results show that EUGENe simplifies the extraction of salient features from models trained within the same workflow.

#### JunD ChIP-seq binding classification

As our final use case and to further demonstrate extensibility, we applied EUGENe to the classification of JunD binding as described in Kopp *et al*^38^. This task utilizes ChIP-seq data from ENCODE^2^ to generate input sequences and binarized classification labels for each sequence (**Figure 5a**). Briefly, regions of interest (ROIs) were defined by extending called peaks by 10,000bp in each direction. These ROIs were then segmented into non-overlapping 200bp bins, with bins that overlap the original peak assigned a positive label and all others assigned a negative label. The genomic sequences represented by each bin were then extended in each direction to increase the context seen as input (150bp in this case for a final input length of 500bp) and used to train neural network classifiers. To first build a DL ready dataset for this prediction task, we wrapped data loading functions from the Janggu package^38^. These functions allow EUGENe users to directly read data from BED, BAM and BigWig file formats into SeqData objects. We then implemented the CNN architecture described in Kopp *et al* (Kopp21CNN) and benchmarked classification performance against built-in FCNs, CNNs, and Hybrid models with matched hyperparameters (**Supplementary Table 9**). All models were configured to incorporate information from both the forward and reverse strand (double stranded or “ds” models).

**Figure 5.**
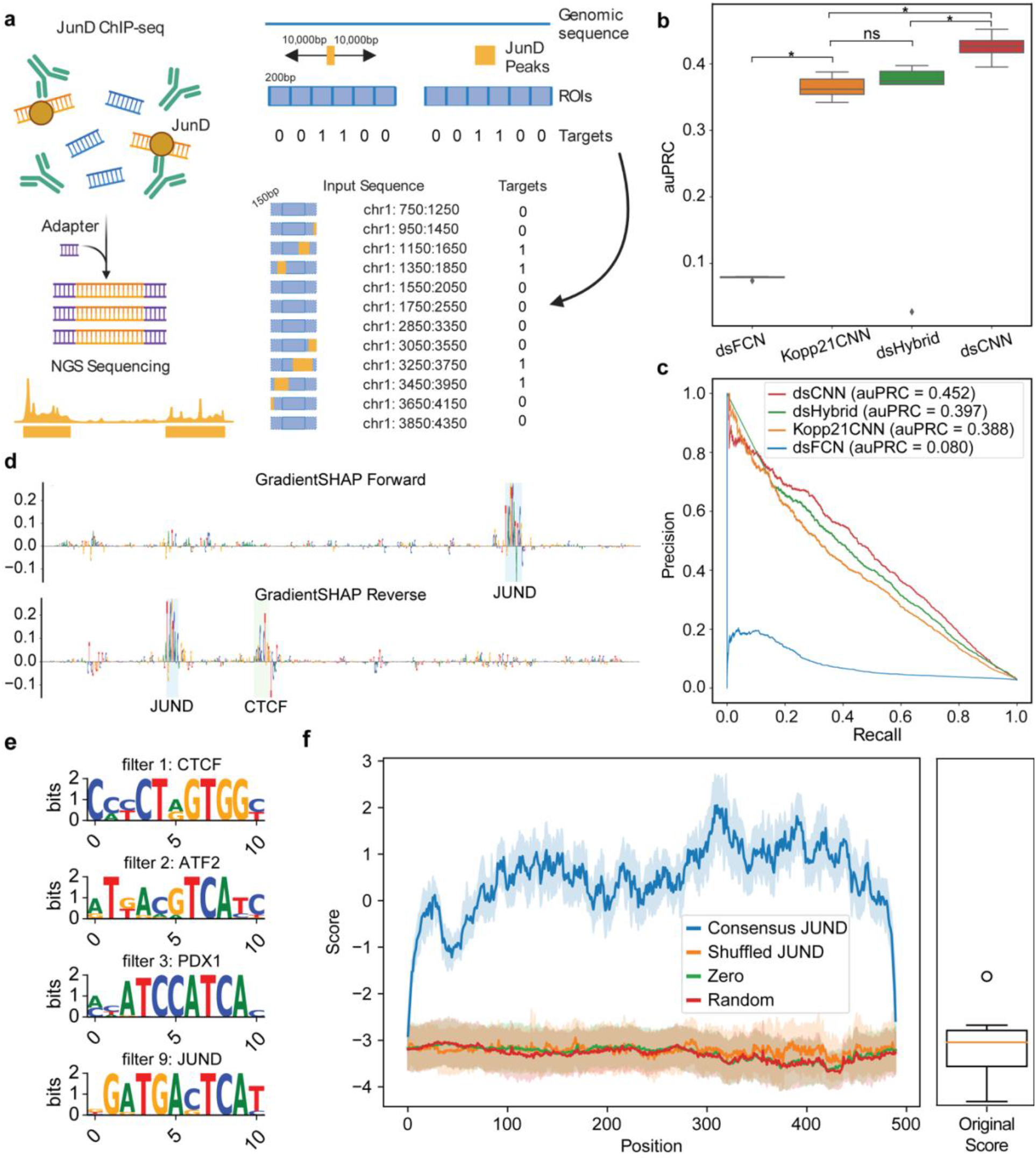
JunD ChIP-seq binding prediction identifies JunD motifs. **a**, kopp21 use case schematic. We used Janggu data loaders to load in a set of 11,086 ChIP-seq peaks for JunD and to generate positive and negative sets for JunD binding prediction. The data loaders take in a set of regions of interest (ROIs) along with peaks and a bin size and output a set of labeled sequences for each bin in the ROI. Bins are labeled as positive (1) if they overlap a peak and negative (0) if they do not. Upon loading, each sequence is extended by 150bp in each direction to provide more sequence context for prediction. **b, c** auPRCs on held-out test data from chromosome 3 for JunD binding classification across four double-stranded architectures **b**, A boxplot across 5 random initializations of each model. **c**, auPR curves for the best models from each architecture. **d**, Feature attribution sequence logos for the top predicted sequence. The top row shows attributions from the forward strand and the bottom row from the reverse strand. Attributions were calculated using GradientSHAP. **e**, A selected set of convolutional filters visualized as PWM logos with significant annotations from TomTom. **f**, Model scores for 10 random sequences with consensus JunD motif implanted at each possible location. 95% confidence intervals of scores are also shown. The boxplot shows the distribution of scores for the random sequences prior to JunD motif implantation. *p < 0.05, ns = not significant. Mann-Whitney U test p-values corrected by the Benjamini-Hochberg method.

We trained models using the same procedure described in Kopp *et al* (see **Methods**)^38^, again with 5 random initializations per architecture to assess reproducibility. Due to the unbalanced nature of the dataset, we focused on evaluating models with the area under the precision recall curve (auPRC). For our Kopp21CNNs, we were able to achieve comparable performances on held out chromosome 3 sequences to those reported by Kopp *et al* for one-hot encoded sequences (**Figure 5bc, Supplementary Table 10**). The dsFCN, the only model without any convolutional layers, immediately overfit the data after a single training epoch and was not at all predictive of binding (**Figure 5c**). The dsCNN models, however, achieved higher auPRCs than both the dsHybrid and Kopp21CNN architectures.

We next applied EUGENe’s interpretation tools to ask whether our best models were learning sequence features relevant to JunD binding to make predictions. We first generated feature attribution scores for the forward and reverse complement strands of all test set sequences using the GradientSHAP method and visualized the most highly predicted sequences as sequence logos (**Figure 5d, Supplementary Figure 5a**). We observed that the most important nucleotides often highlighted consensus or near consensus JunD motifs and that these motifs were often attributed similarly on both the forward and reverse strands (**Figure 5d, Supplementary Figure 5a)**. However, there were instances where a salient motif was highlighted on one strand but not the other (**Figure 5d, Supplementary Figure 5a)**. **Figure 5d** shows one such instance where a CTCF binding site is highlighted on the reverse strand but not the forward strand, indicating the utility of incorporating information from both strands for prediction. We next generated PFM representations for all 10 filters of each convolutional model (excluding dsFCNs) and annotated them using TomTom against the HOCOMOCO database^54^ (**Figure 5e, Supplementary Figure 5b, Supplementary Table 11**). Among the top hits, we found several filters annotated with motifs such as JunD and CTCF (**Figure 5e**, **Supplementary Figure 5b**). Finally, we performed an *in silico* experiment with the best dsCNN model where we slid a consensus JunD motif across each position of a set of 10 randomly generated sequences and predicted binding (**Figure 5f**). We observed that the simple inclusion of the consensus binding site led to a significant jump in predicted output with some position specificity. These results once again showcase that EUGENe’s interpretation methods can help explain model predictions, in this case for DNA protein binding from a genome wide assay.

## Discussion

Despite numerous recent advances and successes in the space, the progress of DL in regulatory genomics has been hindered by the fragmented nature of the set of tools, methods and data that exist across the field. With EUGENe, we seek to integrate many of these aspects into an ecosystem of Python software that provides users with unprecedented functionality. We designed EUGENe to streamline the development of DL workflows in genomics by creating a simplified interface for loading and preparing sequence datasets, instantiating and training neural networks, and evaluating and interpreting trained models. We introduced two data structures that form the basis of EUGENe workflows, SeqData and BaseModel, that allow for both abstraction from and control over the technical details of the workflow. Finally, we demonstrated the versatility of the toolkit by implementing, training and interpreting a variety of regression and classification architectures to model three distinct tasks and datasets.

The single cell field is a particularly attractive guide for developing such an ecosystem. By creating standards for data structures, user interfaces, documentation pages and coding principles, tools such as Scanpy^43^, scVI^61^ and muon^71^ have greatly simplified single cell workflows. We mimicked the structure of many of the tools that sit within the overall single cell software universe by making EUGENe highly modular and wholly contained within the larger Python ecosystem (Pandas, NumPy, scikit-learn, etc.). This gives future contributors the power to easily extend and integrate the functionality currently available in our tool.

There are numerous opportunities for future development of EUGENe, but we see a few as high priority. EUGENe is primarily designed to work on nucleotide sequence input (DNA and RNA), but currently does not have dedicated functions for handling protein sequence or multi-modal inputs. As assays move from bulk to single cell resolution, it will also be important to develop functionality for handling single cell data that allows users to easily ask questions about cell type specific regulatory syntax. Furthermore, we note that SeqData objects currently must be read entirely into memory, which can be a bottleneck for training on very large datasets or with limited compute resources. AnnData^59^ and Janggu^38^ are capable of loading views of data that are never fully stored in memory and we anticipate updating SeqData to behave in a similar manner. We also do not currently offer any dedicated functionality for hyperparameter optimization. Though users familiar with libraries like RayTune^72^ and Optuna^73^ could still utilize many of the objects and functions provided by EUGENe to generate their own hyperoptimization routines, we plan on developing simplified wrappers for performing hyperparameter optimizations (and other training routines) that are native to EUGENe in the future. Similarly, though any user could develop their own methods for benchmarking EUGENe models against shallow machine learning models like gkm-SVMs^74^ or random forests^75^, we plan on integrating functionality for automating this process. Finally, we plan on expanding EUGENe’s dataset, model, metric and interpretation^45–49,51^ library to encompass a larger portion of those available in the field.

As large consortia (such as ENCODE Phase 4 and Impact of Genomic Variation on Function) and individual groups continue to generate functional genomics data at both the bulk and single cell level, the need for a standardized deep learning analysis ecosystem to handle this data becomes even more pressing. We believe that EUGENe represents a positive step in the direction of developing such an ecosystem. Building off this work will allow computational scientists to rapidly develop and share methods and models that answer important questions about the regulatory sequence code.

## Methods

### Analysis of plant promoter data

#### Data acquisition and preprocessing

Plant promoter assay data were obtained from the GitHub repository associated with Jores *et al*. These included two identical libraries for a set of 79,838 plant promoters synthesized with an upstream viral 35S enhancer and downstream barcode tagged GFP reporter gene (**Figure 3a**). The libraries were designed to include 10-20 constructs with distinct barcodes for each promoter. These libraries were used to transiently transform both tobacco leaves and maize protoplasts and promoter activities were assayed using plant STARR-seq^63^. Per barcode activity was calculated as the ratio of RNA barcode frequency to DNA barcode frequency and the median of these ratios was then used to aggregate across barcodes assigned to the same promoter. These aggregated scores were then normalized by the median value for a control construct and were log transformed to calculate a per promoter “enrichment” score. We downloaded these enrichment scores (https://github.com/tobjores/Synthetic-Promoter-Designs-Enabled-by-a-Comprehensive-Analysis-of-Plant-Core-Promoters/tree/main/CNN) for both libraries as separate datasets which we could use as training targets. We used the identical 90/10 training and test split used in Jores *et al* (the dataset could be downloaded with set labels). The training set was further split into 90/10 train and validation sets. All sequences were one-hot encoded using a channel for each letter of the DNA alphabet (“ACGT”).

#### Model initialization and training

We implemented the Jores21CNN architecture by translating the Keras code in the associated GitHub repository into PyTorch and integrating it into our library. We benchmarked this architecture against built-in CNN and Hybrid architectures in EUGENe with the hyperparameters described in **Supplementary Table 3**. In each convolutional layer, the Jores21CNN first applies a set of filters to the input as is standard for convolutional models, but also applies the reverse complements of the filters (as opposed to the reverse complement of the sequences) to each input in an effort to capture information from both strands^64^. Since this still only requires a single strand as input into the models, we opted to benchmark against only single stranded (ss) versions of built-in CNN and Hybrid models. Following instantiation, we initialized 78 filters in the first convolutional layer of each model using PWMs derived from core promoter elements and transcription factor binding clusters downloaded from the GitHub repository (https://github.com/tobjores/Synthetic-Promoter-Designs-Enabled-by-a-Comprehensive-Analysis-of-Plant-Core-Promoters/tree/main/data/misc) associated with the publication. All other parameters were initialized by sampling from the Kaiming normal distribution^69^. We trained models for a maximum of 25 epochs with a batch size of 128 and used the Adam optimizer with a starting learning rate of 0.001. We also included a learning rate scheduler that modified the learning rate during training with a patience of 2 epochs. We used mean squared error as our objective function and stopped training early if the validation set error did not decrease after 5 epochs.

#### Model evaluation and interpretation

Models were primarily evaluated using the percentage of variance explained (R^2^) on predictions for the test set. We repeated the above training procedure across 5 independent random initializations and evaluated R^2^ scores across these trials. For PWM visualization, we used the approach described in Minnoye *et al*^17^. Briefly, for each filter in the first convolutional layer, we calculated activations for all subsequences (of the same length as the filter) within the test set sequences. We then took the top 100 subsequences corresponding to the top 100 activations (maximally activating subsequences) and generated a PFM. For visualizing filters as sequence logos, we converted PFMs to PWMs using a uniform background nucleotide frequency. We calculated feature attributions for all test set sequences using the DeepLIFT method. To perform the feature implantation approach, we downloaded the 16bp PFM containing the consensus TATA box motif from the Jores *et al* GitHub repository and one-hot encoded it by taking the highest probability nucleotide at each position. We also downloaded the set of 310 promoters (https://github.com/tobjores/Synthetic-Promoter-Designs-Enabled-by-a-Comprehensive-Analysis-of-Plant-Core-Promoters/blob/main/analysis/validation_sequences/promoters_for_evolution.tsv) used in Jores *et al* for *in silico* evolution. We then implanted the TATA box containing sequence at every possible position of each of the 310 promoter sequences and used the best performing models (one each from leaf, protoplast and combined) to make predictions. We compared this to predicted scores generated with the same feature implantation approach using a dinucleotide shuffled version of the 16bp sequence containing the TATA box motif, a random 16bp one-hot encoded sequence, and a 16bp all zeros input. We performed the *in silico* evolution experiments on the same set of 310 promoter sequences^31,62^. In each round, we first used *in silico* saturation mutagenesis to identify the mutation that increased the model score by the largest positive value (delta score). We then introduced this mutation into the sequence and repeated this for 10 iterations.

### Analysis of RNA binding data

#### Data acquisition and preprocessing

As described in detail in Alipanahi *et al*, a set of 241,357 31-41nt long RNA probes were split into two experimental sets, Set A and Set B, with each designed to include all possible 9-mers at least eight times, all possible 8-mers at least 33 times and all possible 7-mers 155 times **(Figure 4a**). These probes were assayed against 244 RBPs using a protein binding microarray (PBM) ^68^, and intensities were normalized as described in Ray *et al*^67^. We downloaded the normalized RNA probe binding intensity matrix from the Ray *et al* supplementary information (http://hugheslab.ccbr.utoronto.ca/supplementary-data/RNAcompete_eukarya/norm_data.txt.gz) and separated the Set A and Set B sequences into two distinct groups. To remove outliers, we set all values of probe intensities to be capped at the 99.95 percentile for each prediction task (RBP). We then Z-scored the clamped values to zero mean and unit standard deviation for each RBP. All normalizations were performed using Set A statistics (i.e. Set B values were z-scored using means and standard deviations from Set A). For multitask prediction, we removed the 11 RBPs with ≥ 0.1% missing values across all probes in Set A, and further removed all probes in Set A that had any missing values for any of the remaining 233 RBPs. This left 120,326 and 110,645 probes for training single task and multitask models respectively and 121,031 in Set B for testing. Set A was then further split 80/20 into a training and validation set. All sequences were one-hot encoded using a channel for each of the RNA alphabet (“ACGU”) for input into models.

#### Model initialization and training

We implemented the DeepBind architecture (https://github.com/cartercompbio/EUGENe/blob/main/eugene/models/_sota_models.py) described in the Supplementary Information (https://static-content.springer.com/esm/art%3A10.1038%2Fnbt.3300/MediaObjects/41587_2015_BFnbt3300_MOESM51_ESM.pdf) of Alipanahi *et al* and added it as a EUGENe SOTA model. DeepBind architectures were initially designed to take either the forward strand (ss) or both strands (ds) as input. However, Alipanahi *et al* trained their RBP models with just the single strand input due to the single stranded nature of RNA, so we also used a single stranded (ss) implementation for our DeepBind models. We initialized both the single task models and the multitask model with parameters sampled from the Kaiming normal distribution^69^ and trained all models for a maximum of 25 and 100 epochs respectively, using the Adam optimizer^70^ and a starting learning rate of 0.005. We also included a learning rate scheduler that modified the learning rate during training with a patience of 2 epochs. The batch size for training was fixed to 64 and 1024 for single- and multi-task models respectively and mean squared error was used as the objective function for all models, with training halting if the validation set error did not decrease after 5 epochs. For multitask models, we used the average mean squared error across all tasks. Hyperparameters selected for the architectures of each model are provided in **Supplementary Table 6**. Finally, we downloaded a set of 89 pretrained human RBP models (https://kipoi.org/models/DeepBind/Homo_sapiens/RBP/) from Kipoi and wrapped functions from the Kipoi package to make predictions using these models.

#### Model evaluation

We evaluated models using the Z-score, AUC and E-score metrics reported in Alipanahi *et al*. To calculate these metrics, we first computed a binary *n x m* matrix *A*, where the *n* rows represent all possible 7-mers from the RNA alphabet (AAAAAAA, AAAAAAC, AAAAAAG, etc.) and the *m* columns represent the 121,031 probes assayed from Set B. Each entry *a_ij_* in the matrix is 1 if the *i*th k-mer is found in the *j*th probe, and 0 otherwise. Consider first working with a single RBP, in which we have normalized binding intensity values for each of the 121,031 probes (*m*-dimensional vector *x*). We compared the *i*th row (representing a k-mer) of the matrix *A* (an *m*-dimensional vector) to the vector *x* of observed intensities and computed the Z-scores, AUCs and E-scores for that k-mer as described in ^9^ and ^67^. We repeated this for all k-mers (across rows of *A*) to generate an *n* dimensional vector for each metric meant to capture the importance of each k-mer for binding that RBP. For Z-scores, 0 indicates an average level of binding when that k-mer is present in the probe sequence, with more positive scores indicating higher levels of binding than average when that k-mer is present. For AUC and E-scores (a modified AUC), the value is bound between 0 and 1, with values closer to 1 indicating more binding when that k-mer is present. We repeated this process for all models that predict probe intensities by substituting the predicted intensities from a given model for the vector *x* of observed intensities. We generated a set of *n* dimensional vectors for each model-metric pair (i.e. for a single task model, we have a vector for Z-scores, a vector for E-scores and a vector for AUCs) then took each of these vectors and calculated Pearson and Spearman correlations with the vector *x* from the observed Set B intensities. This results in a pair of correlation values, one Pearson and one Spearman, describing the performance of a given model on a specific RBP (these are single points in the boxplots shown in **Figure 4b and Supplementary Figure 4a**).Repeating this process for all RBPs generates a distribution of correlations for a given model.

We can use the same procedure on Set A observed intensities to generate a distribution of correlations analogous to a biological replicate. These are the “Set A” and “Observed intensities” columns of **Figure 4b and Supplementary Figure 4a**. We generated the distribution labeled “Set A” in this way with our own implementation of these metrics and downloaded the “Observed intensities” distribution from performance tables included in the supplement of Alipanahi *et al*. Finally, we calculated Pearson and Spearman correlation coefficients for the observed and predicted intensities on Set B for all models. Note that this is not possible to do for Set A since the probes are different for this set, hence the omission of the “Set A” and “Observed intensities” columns in the last boxplots of **Figure 4b** and **Supplementary Figure 4a**.

#### Model interpretation

For filter visualization, we used the approach described in Alipanahi *et al*. Briefly, for a given filter, we calculated the activation scores for all possible subsequences (of the same length as the filter) from Set B probes and identified the maximum value. We then used only the subsequences with an activation at least 3/4ths of this maximum to generate a PFM for that filter. We repeated this process for all 32 filters in each of the top 10 single task models and for all 1024 filters of the multitask model. The top 10 single task models were chosen based on ranking of Pearson correlations between observed and predicted intensity values. We then input all multitask PFMs to TomTom for annotation against the Ray *et al* database and filtered for hits with a Bonferroni multiple test corrected p-value ≤ 0.05. We calculated feature attributions for all Set B probes using the InputXGradient method. For multitask models, feature attributions can be calculated on a per task basis to determine how each nucleotide of the input sequence influenced that particular task. We again only did this for a subset of RBPs, using the Pearson correlation of predicted and observed intensities to choose the top 10 single task models and the top 10 predicted tasks for the multitask model. We use the same *in silico* evolution method for this use case as we did for the plant promoters. Using trained models for selected RBPs, we first performed 5 rounds of evolution on 10 randomly generated sequences of 41nt in length (“ACGT” sampled uniformly). We then calculated feature attributes for the initial random sequences and the evolved sequences using the InputXGradient method and compared them.

### Analysis of JunD binding data

#### Data acquisition and preprocessing

We followed the same procedure to acquire and preprocess the data for training models on the prediction of JunD binding as reported in Kopp *et al*^38^. We started by downloading JunD peaks from human embryonic stem cells (H1-hesc) called with the hg38 reference genome from encodeproject.org (ENCFF446WOD, conservative IDR thresholded peaks, narrowPeak format). We next defined regions of interest (ROIs) by extending the union of all JunD peaks by 10kb in each direction. We removed blacklisted regions for hg38 (http://mitra.stanford.edu/kundaje/akundaje/release/blacklists/hg38-human/hg38.blacklist.bed.gz) using bedtools^76^ and trimmed the ends of resulting regions to be divisible into 200bp bins. For training and testing, we binned ROI’s into 200bp sequences and labeled any of those that overlapped a JunD binding peak with a positive label and all non-overlapping bins with a negative label. As input to models, we first extended each genomic bin by 150bp on each side (so that the model sees 500bp in total for each input when predicting on a 200bp site) and then one-hot encoded using a channel for each of the DNA alphabet (“ACGT”). In total, we used 1,013,080 200bp bins for generating training, validation and test sets. We split the sequences by chromosome so that validation sequences were from chr2 and test sequences from chr3 (the rest were used for training).

#### Model initialization and training

For the JunD binding task, we first implemented the Kopp21CNN arhitecture decscribed in Kopp *et al* by following the Keras code in the associated GitHub repository along with their description of the layers in the Supplementary Information (https://static-content.springer.com/esm/art%3A10.1038%2Fs41467-020-17155-y/MediaObjects/41467_2020_17155_MOESM1_ESM.pdf). We then trained 5 random initializations of dsFCNs, dsCNNs, dsHybrids and Kopp21CNNs, each with parameters sampled from the Kaiming normal distribution. All models used both the forward and reverse strands as input through the same architecture (ds). Following Kopp *et al*, we trained all models for a maximum of 30 epochs with the AMSGrad optimizer^77^ and a starting learning rate of 0.001. The batch size for training was fixed to 64 for all models and binary cross entropy was used as the objective function, halting training if the validation set error did not decrease after 5 epochs. Hyperparameters selected for the architectures of each model are provided in **Supplementary Table 9**.

#### Model evaluation and interpretation

Models were primarily evaluated using the area under the precision recall curve (auPRC) as the dataset was heavily imbalanced. We again performed model interpretation using feature attributions, filter visualizations and *in silico* experimentation methods from EUGENe. We calculated feature attributions for the forward and reverse strands of all test set sequences using the GradientSHAP method. To visualize filters, we applied the approach from ^17^ and generated PFMs. We fed these PFMs to the TomTom webserver and queried the HOCOMOCO CORE database^54^. We subset filters down to those with a multiple test corrected p-value ≤ 0.05 and manually inspected the top hits. These PWMs were visualized as logos using a uniform background of nucleotide frequencies. We performed the *in silico* implantation experiment using the JunD PFM downloaded from JASPAR (https://jaspar.genereg.net/matrix/MA0491.1). We calculated model scores by generating 10 randomly generated sequences (uniformly sampled) and implanting the consensus one-hot encoded JunD motif at every possible position. We compared this to predicted scores from applying the same approach to a random one-hot encoded sequence, an all zeros input and a dinucleotide shuffled JunD motif, all of the same length as the consensus JunD motif.

#### Data visualization software

For most exploratory data analysis and performance evaluations, we used a combination of the Seaborn and Matplotlib plotting libraries in Python. For sequence logo visualizations of filters and feature attributions, we used modified functions from the viz_sequence package (https://github.com/kundajelab/vizsequence), the seqlogo package (https://github.com/betteridiot/seqlogo), the seqlogo package (https://github.com/betteridiot/seqlogo) and the logomaker package (https://github.com/jbkinney/logomaker).

## Statistical methods

Mann-Whitney U tests^78^ were used to compare performance distributions between architecture types and p-values were corrected with the Benjamini-Hochberg method^79^. TomTom reports significance of alignments of query motifs to a database using the methods described in ^52^. We used the q-value reported by the webserver tool (https://meme-suite.org/meme/tools/tomtom) and considered hits to be those alignments with a q-value ≤ 0.05 as significant. Figures for *in silico* implantation of motifs included 95% confidence intervals.

## Dataset availability

All datasets used in this study are publicly available. We have collected the specific dataset files and trained models used in the analyses presented here at the following Zenodo link: https://doi.org/10.5281/zenodo.7140082. These represent the raw and processed data files that can be loaded into the EUGENe API to generate the figures in this work using the code linked below. We also have included processed SeqData objects that can be used along with the accompanying code to generate the figures for all use cases.

## Code availability

EUGENe is freely available under the MITlicense at https://github.com/cartercompbio/EUGENe. Documentation for the tool is available at https://eugene-tools.readthedocs.io/en/latest/index.html. Jupyter notebooks and Python scripts used to perform the analyses presented in the use case section are available on GitHub at https://github.com/adamklie/EUGENe_paper.

## Acknowledgements

This work was supported by the National Institutes of Health [grant number 1U01HG012059]; infrastructure was funded by the National Institutes of Health [grant number 2P41GM103504-11]; T.J. is supported by the German Research Foundation [DFG; fellowship number 441540116]. E.K.F and J.J.S were supported by the National Institutes of Health [grant number DP2HG010013]. H.C. is supported by the Canadian Institute for Advanced Research [award number FL-000655]. We would like to thank the community of genomics researchers who made their code open source so that we could utilize it for EUGENe functions. Specifically, we would like to thank the developers of the concise package (https://github.com/gagneurlab/concise) for functions which we modified for the preprocess module, the developers of the deeplift package (https://github.com/kundajelab/deeplift) for functions we modified for dinucleotide shuffling, the developers of the yuzu package (https://github.com/kundajelab/yuzu) for functions we modified for *in silico* saturation mutagenesis, Travis Wrightsman for functions we modified for reading MEME files and the developers of the ExplaiNN (https://github.com/wassermanlab/ExplaiNN) package for functions we modified for saving to MEME.

## Author information

### Authors and Affiliations

Department of Medicine, University of California San Diego, La Jolla, CA, USA

Adam Klie, Joe J. Solvason, Emma K. Farley & Hannah Carter

Bioinformatics and Systems Biology Program, University of California San Diego, La Jolla, CA, USA

Adam Klie, Joe J. Solvason, Emma K. Farley & Hannah Carter

Daniel Hand High School, Madison, CT, USA

Hayden Stites

Department of Genome Sciences, University of Washington, Seattle, WA, USA

Tobias Jores

Department of Biological Sciences, University of California San Diego, La Jolla, CA, USA

Joe J. Solvason & Emma K. Farley

### Contributions

A.K., J.J.S., E.K.F. and H.C. designed the study. A.K. designed the toolkit. A.K. and H.S. implemented the code. A.K. performed the use case analyses. J.J.S. and H.S. performed software testing. H.C. supervised the work. All authors read and corrected the final manuscript.

### Corresponding authors

Correspondence to Hannah Carter.

## Supplementary information

**Supplementary Figure 1.**
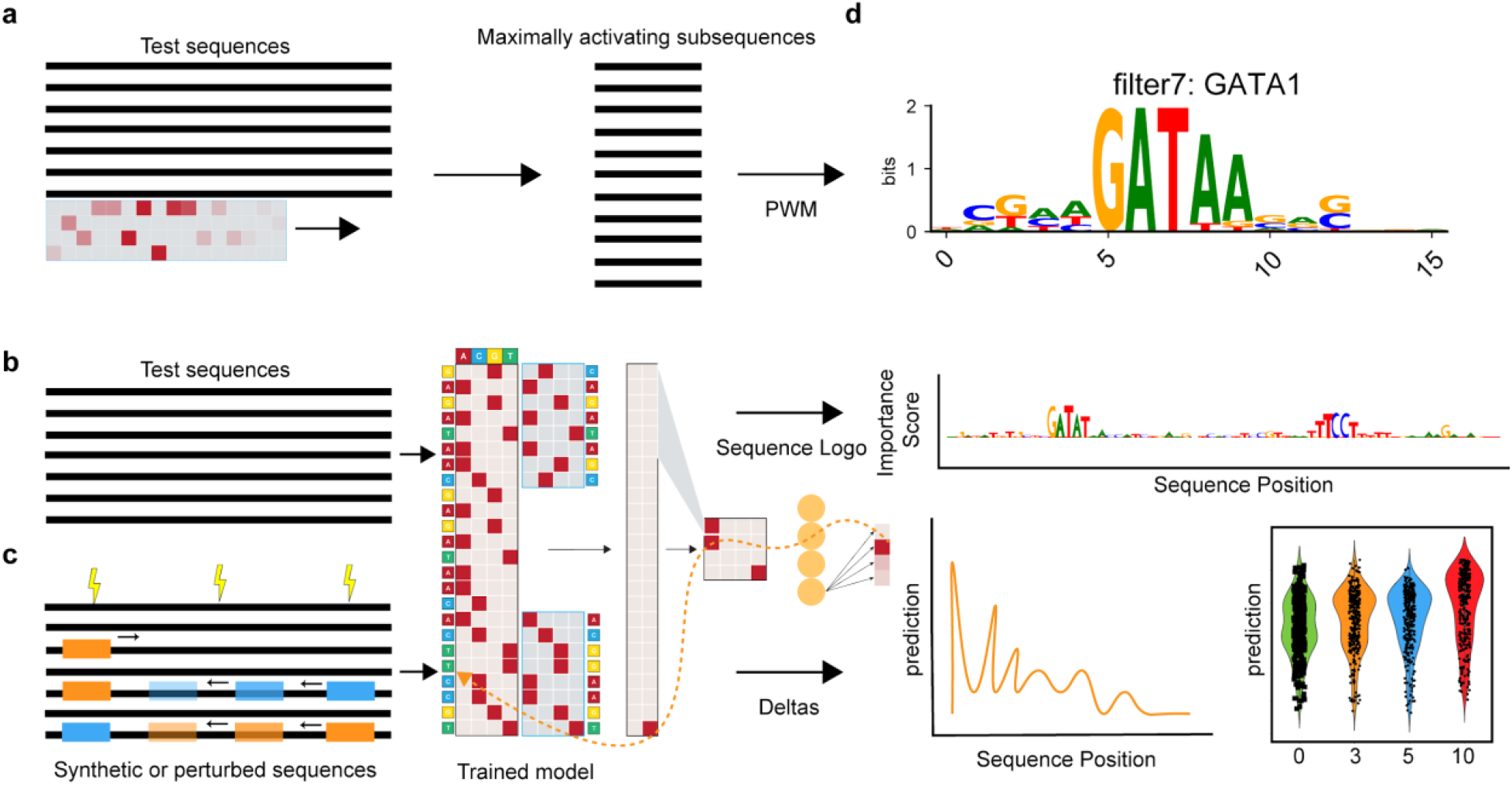
Interpretation methods implemented and visualized in EUGENe. The starting point for most of the interpretation methods in EUGENe are a set of sequences. **a**, For PWM visualization, each filter in the first convolutional layer of a given model is used to scan this set of input sequences to search for filter length subsequences that highly activate the filter. These “maximally activating subsequences” are then used to generate a position frequency matrix that can be transformed to a position weight matrix and visualized as a logo**. b**, We implement several gradient based feature attribution approaches in which sequences are first passed through the model to generate an output. This output signal is then backpropogated through the model parameters back to the input to generate a per nucleotide score that can also be visualized as a sequence logo. **c**, We also provide users functions for running *in silico* experiments using the model as an oracle. Random or synthetically designed sequences that have been mutated or have had motifs implanted in them can be scored using a trained model. The difference in scores between the original sequence or a reference sequence can also be calculated to prioritize functional mutations or feature dependencies that can be experimentally validated. **d**, The methods illustrated in **a, b**, and **c** generate results that can be visualized by function calls on SeqData objects. Toy examples of these visualizations are shown in **d**.

**Supplementary Figure 2.**
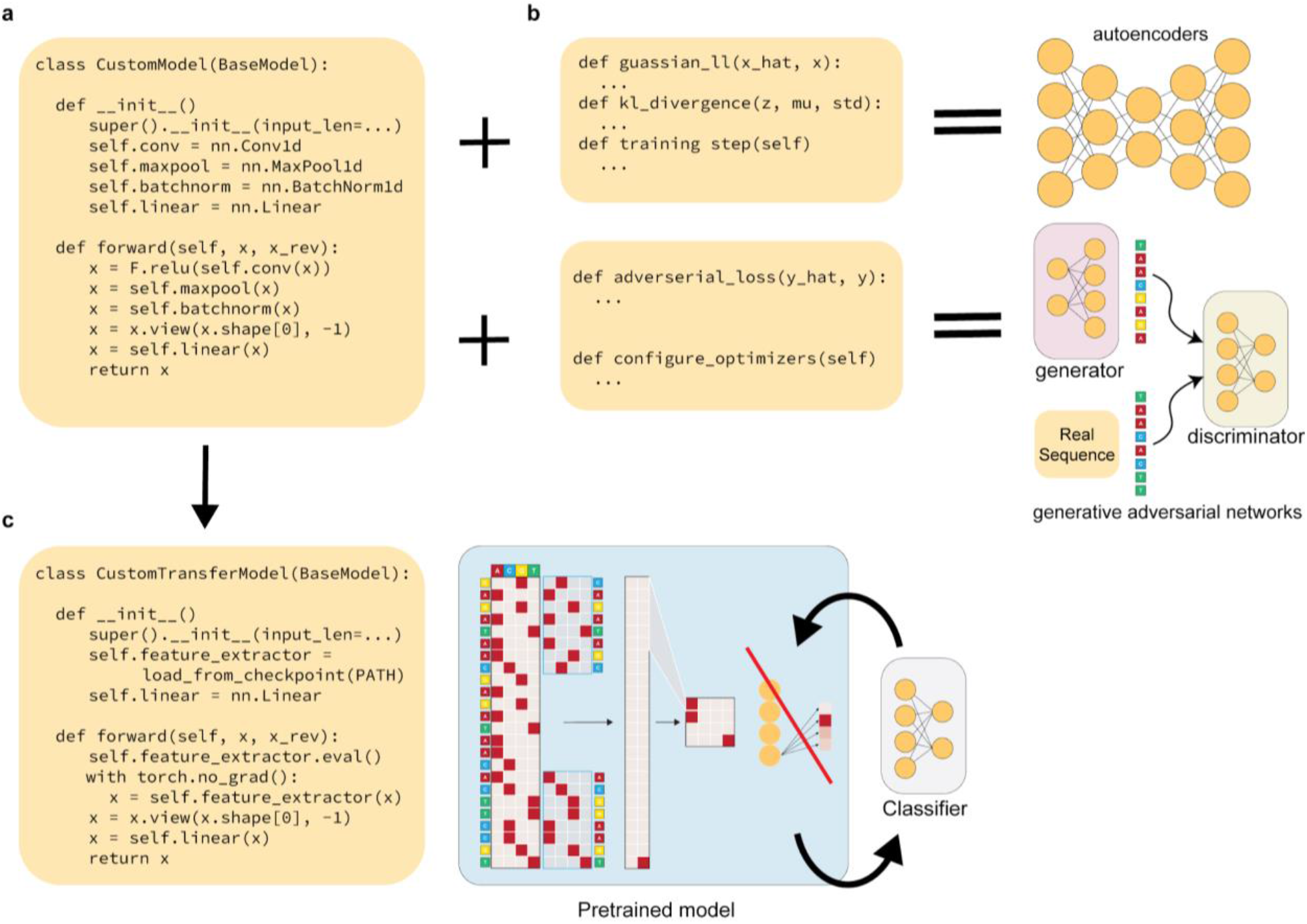
Extending EUGENe’s BaseModel to implement custom architectures. **a**, Creating custom models that are compatible with EUGENe’s basic training protocol involves first inheriting from the BaseModel class (not shown), then defining the model’s architecture (__init__) and the forward propagation (forward) method. **b**, The BaseModel class can also be extended to create variational autoencoders (VAEs) or generative adversarial networks (GANs). A VAE (in its most basic form) requires creating two functions for calculating different parts of the loss and implementing how the functions are integrated into the training function. We have omitted the changes needed to define an encoder and decoder structure and how that is handled in forward. A GAN requires implementing a multipart loss function and configuring multiple optimizers to handle the training of the generator and discriminator. **c**, Transfer learning from pretrained models can be accomplished with simple changes to the initialization and forward functions. Namely, a pretrained PyTorch Lightning model needs to be loaded in the __init__ method and then utilized in the forward method.

**Supplementary Figure 3.**
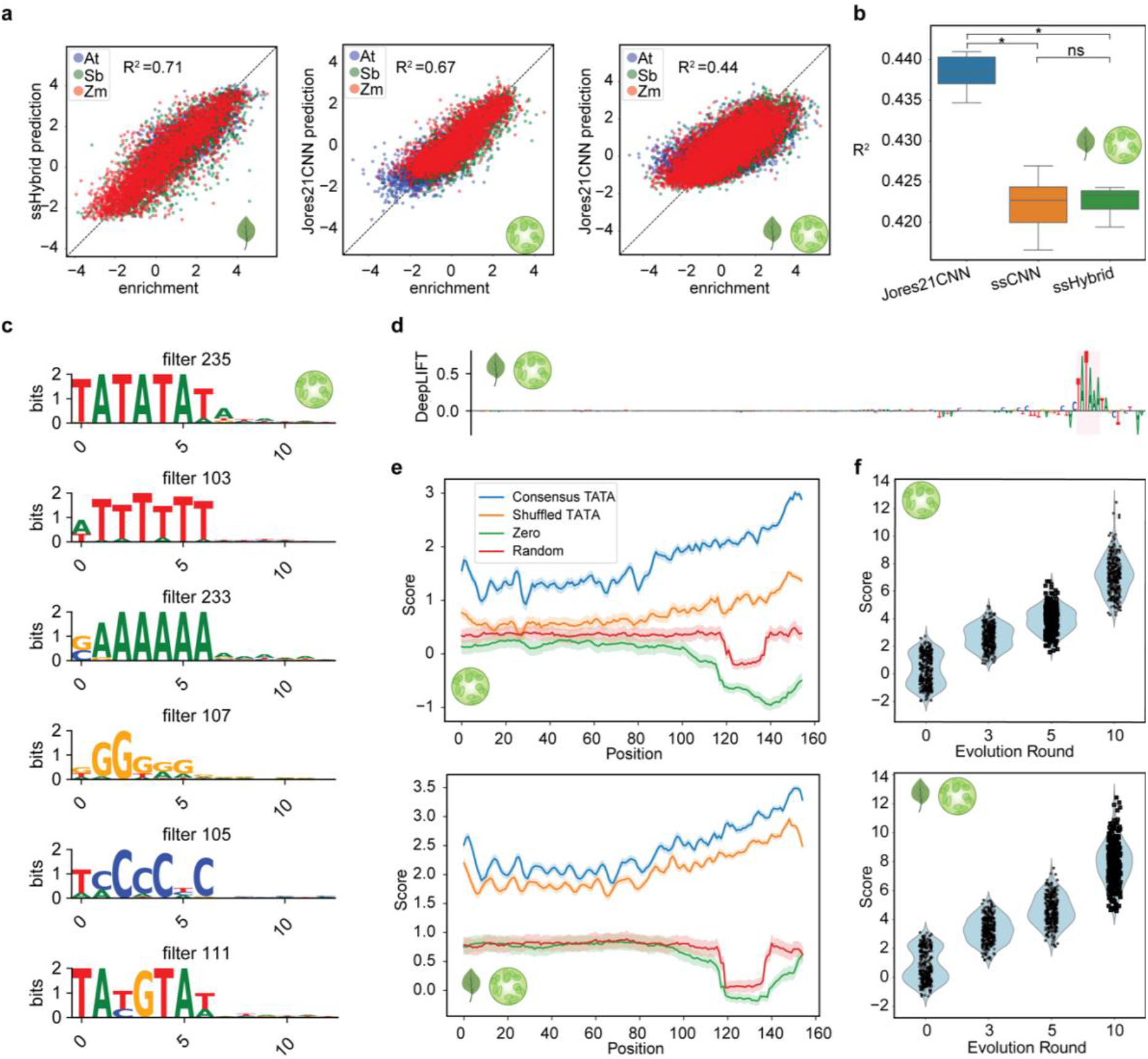
STARR-seq plant promoter activity prediction. **a**, Performance scatterplots colored by species of origin for the best leaf (left), protoplast (middle) and combined (right) models. **b**, Predictive performance of all trained combined models. The boxplots show distributions of R^2^ values on held-out test data for each architecture across 5 random initializations. **c**, PWMs for a hand-selected set of learned protoplast model filters (not initialized with known PWMs). **d**, Feature attribution scores calculated using the DeepLIFT method for the sequence with the highest predicted value in the best combined model **e**, Best protoplast (top) and combined (bottom) model scores for 310 sequences with an implanted consensus TATA box motif, shuffled consensus TATA box motif, all zeros motif, and random motif at every possible position. The 95% confidence interval is shown. **f**, Model scores for the same set of 310 promoters at different rounds of evolution compared against baseline (0) for the best protoplast (top) and combined (bottom) model.

**Supplementary Figure 4.**
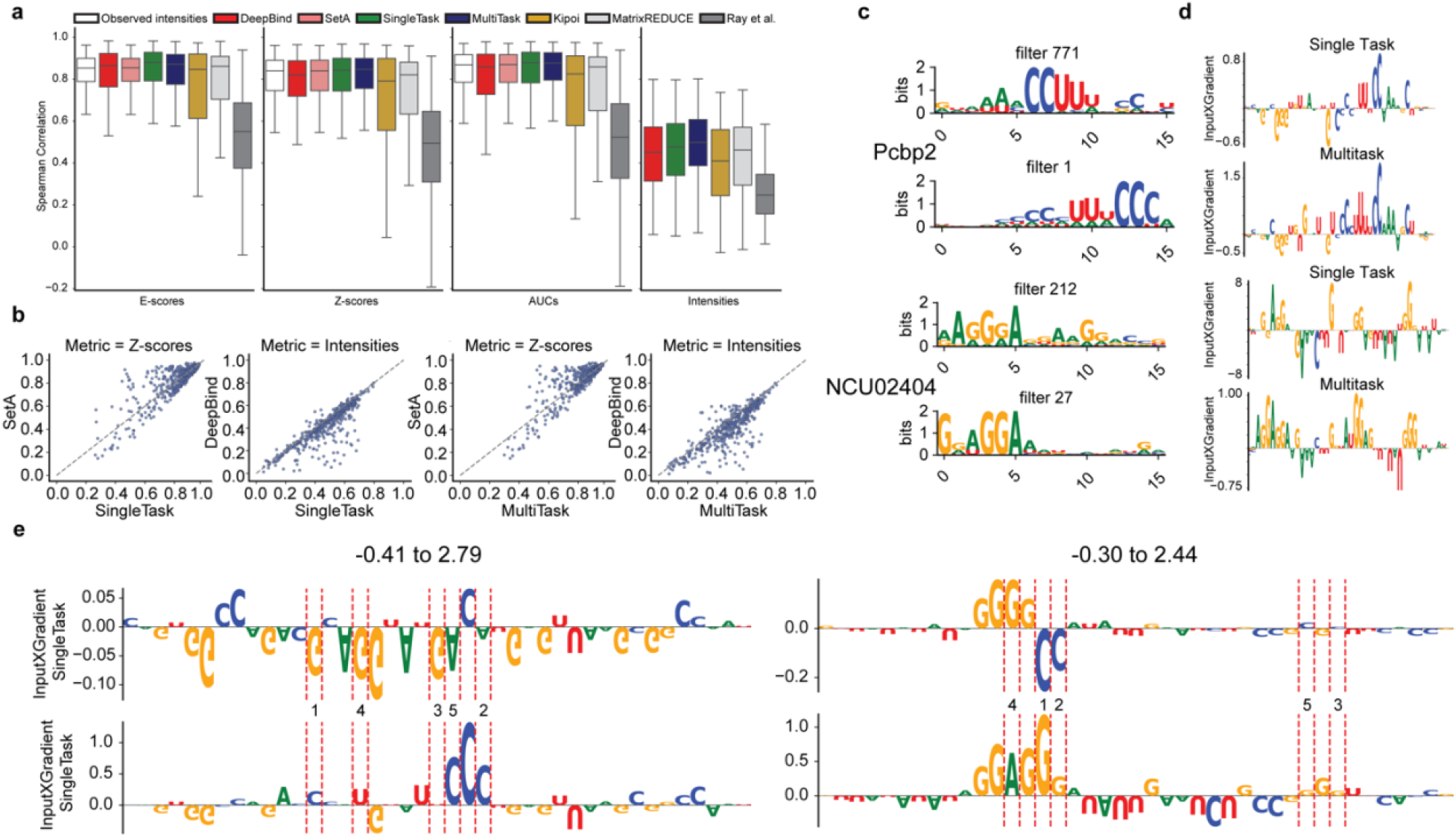
RNA binding protein (RBP) specificity prediction. **a**, Spearman correlations across four different metrics with each metric calculated from comparisons between observed (Set B) and predicted binding intensities (see **Methods** for more details on how each metric is calculated). Each boxplot indicates a distribution of Spearman correlations across all 244 RBPs. Ray et al, MatrixREDUCE, DeepBind and Observed intensities refer to correlations calculated from predicted intensities reported in Alipanahi *et al*. Observed intensities and SetA refer to correlations calculated using the intensities from Set A probes as the predicted intensities (see **Methods**). **b**, Performance comparison scatterplots for the indicated models and metrics. Each dot indicates a comparison of the Pearson correlation between two models on a single RBP. **c**, Multitask and single task filters with TomTom significant annotations for Pcbp2 (top) and NCU02404 (bottom). **d**, Feature attributions calculated using the InputXGradient method for single task and multitask models using the sequence with the highest observed intensity in the test set for Pcbp2 (top) and NCU02404 (bottom). **e**, Two more examples of InputXGradient attribution scores for random (top row) and evolved (bottom row) sequences after evolution with the Pcbp2 (left) and NCU02404 (right) single task models. Red dashed lines indicate mutations made during evolution annotated with the round the mutation occurred in.

**Supplementary Figure 5.**
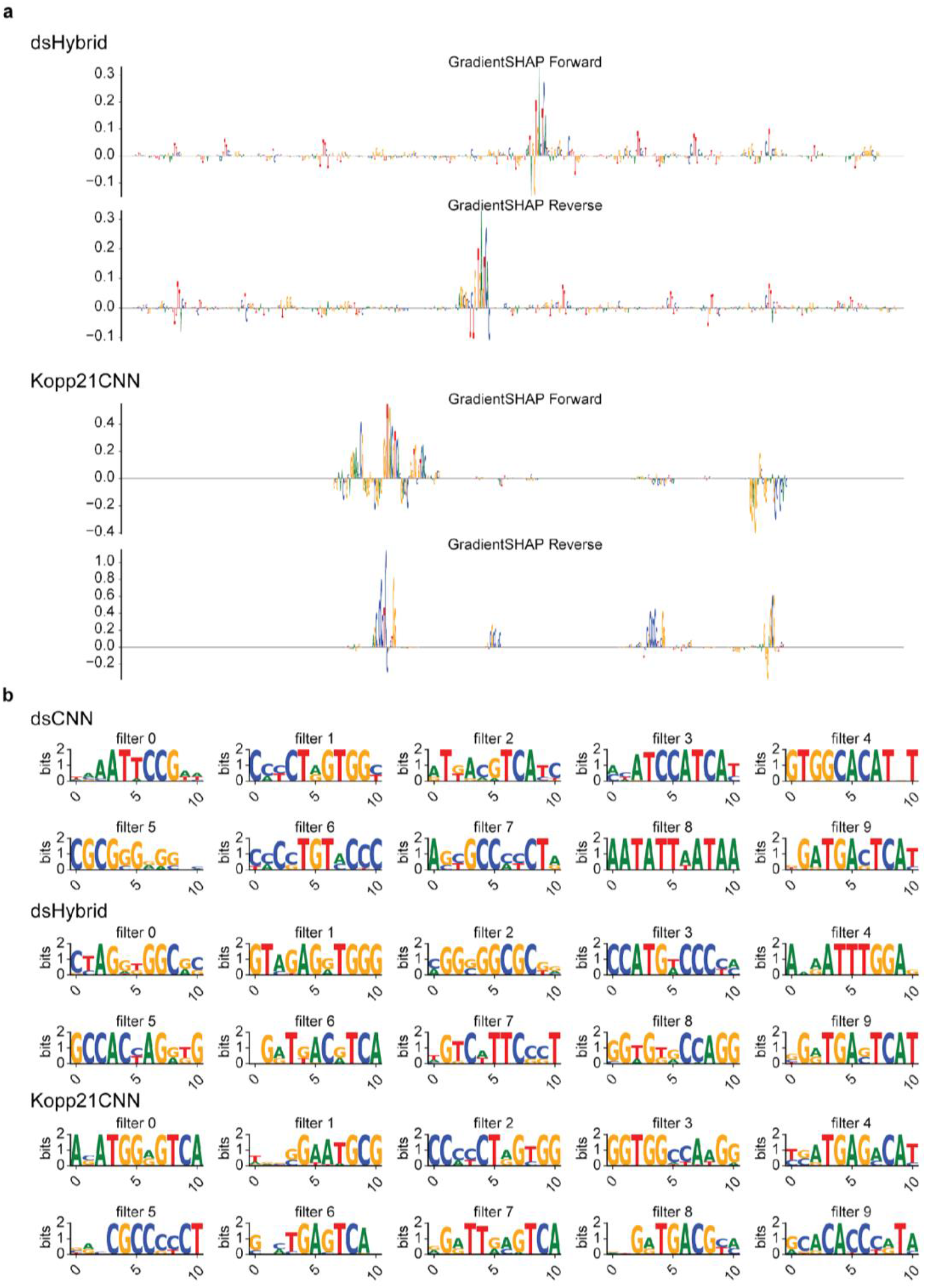
JunD binding classifier interpretation. **a**, Feature attribution scores calculated using GradientSHAP for the forward and reverse complement of the sequence with the highest predictions in each of the dsHybrid and Kopp21CNN models. **b**, PWM visualizations of the 10 filters for the three convolutional architectures trained for JunD binding classification.

All supplementary tables can be found in supplementary_tables.xlsx.

