## Supplementary Figure for "EUGENe: A Python toolkit for predictive analyses of regulatory sequences"

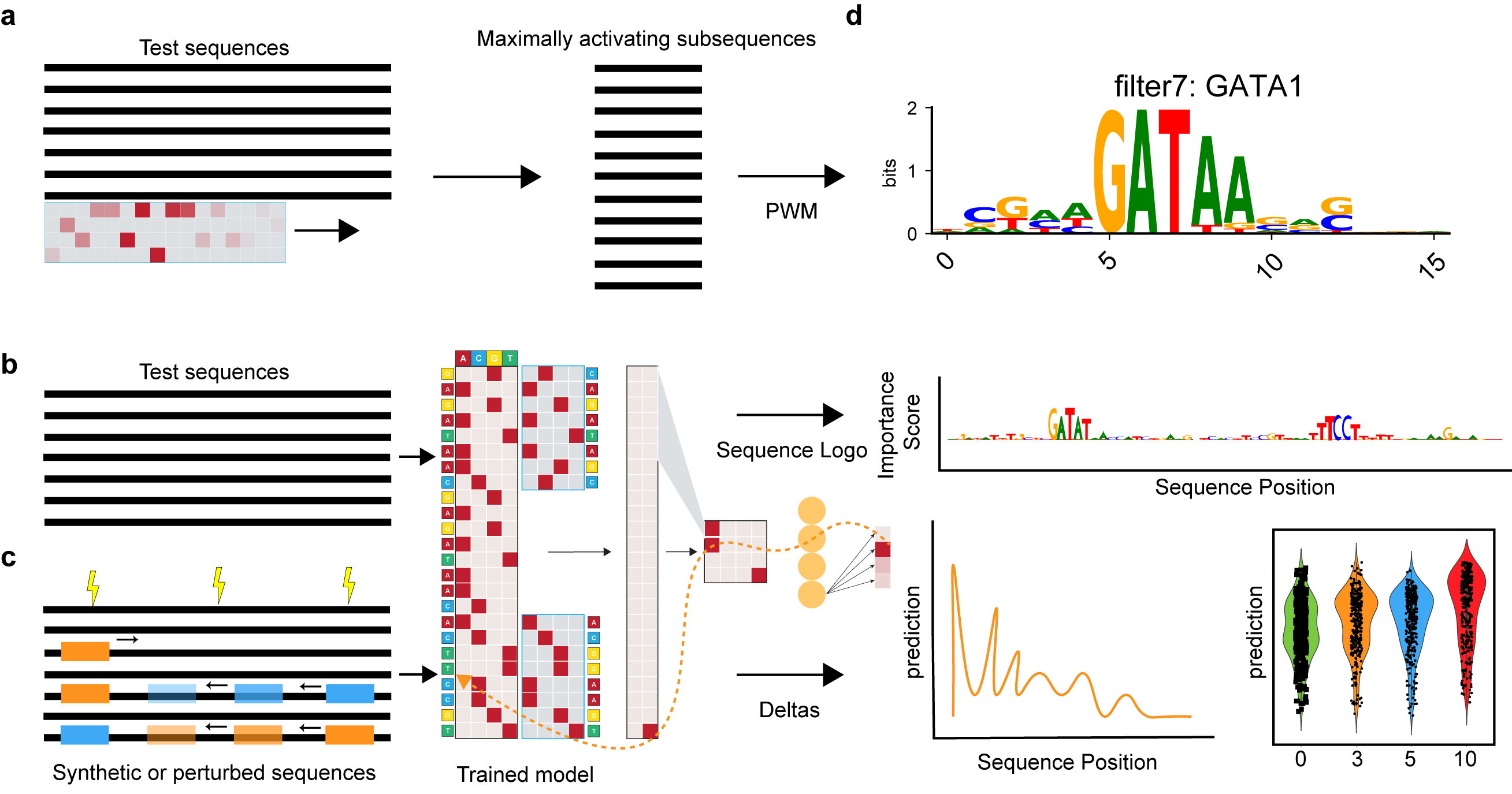

**Supplementary Figure 1. Interpretation methods implemented and visualized in EUGENe**. The starting point for most of the interpretation methods in EUGENe are a set of sequences. **a,** For PWM visualization, each filter in the first convolutional layer of a given model is used to scan this set of input sequences to search for filter length subsequences that highly activate the filter. These “maximally activating subsequences” are then used to generate a position frequency matrix that can be transformed to a position weight matrix and visualized as a logo**.** **b,** We implement several gradient based feature attribution approaches in which sequences are first passed through the model to generate an output. This output signal is then backpropogated through the model parameters back to the input to generate a per nucleotide score that can also be visualized as a sequence logo. **c,** We also provide users functions for running *in silico* experiments using the model as an oracle. Random or synthetically designed sequences that have been mutated or have had motifs implanted in them can be scored using a trained model. The difference in scores between the original sequence or a reference sequence can also be calculated to prioritize functional mutations or feature dependencies that can be experimentally validated. **d,** The methods illustrated in **a, b,** and **c** generate results that can be visualized by function calls on SeqData objects. Toy examples of these visualizations are shown in **d**.

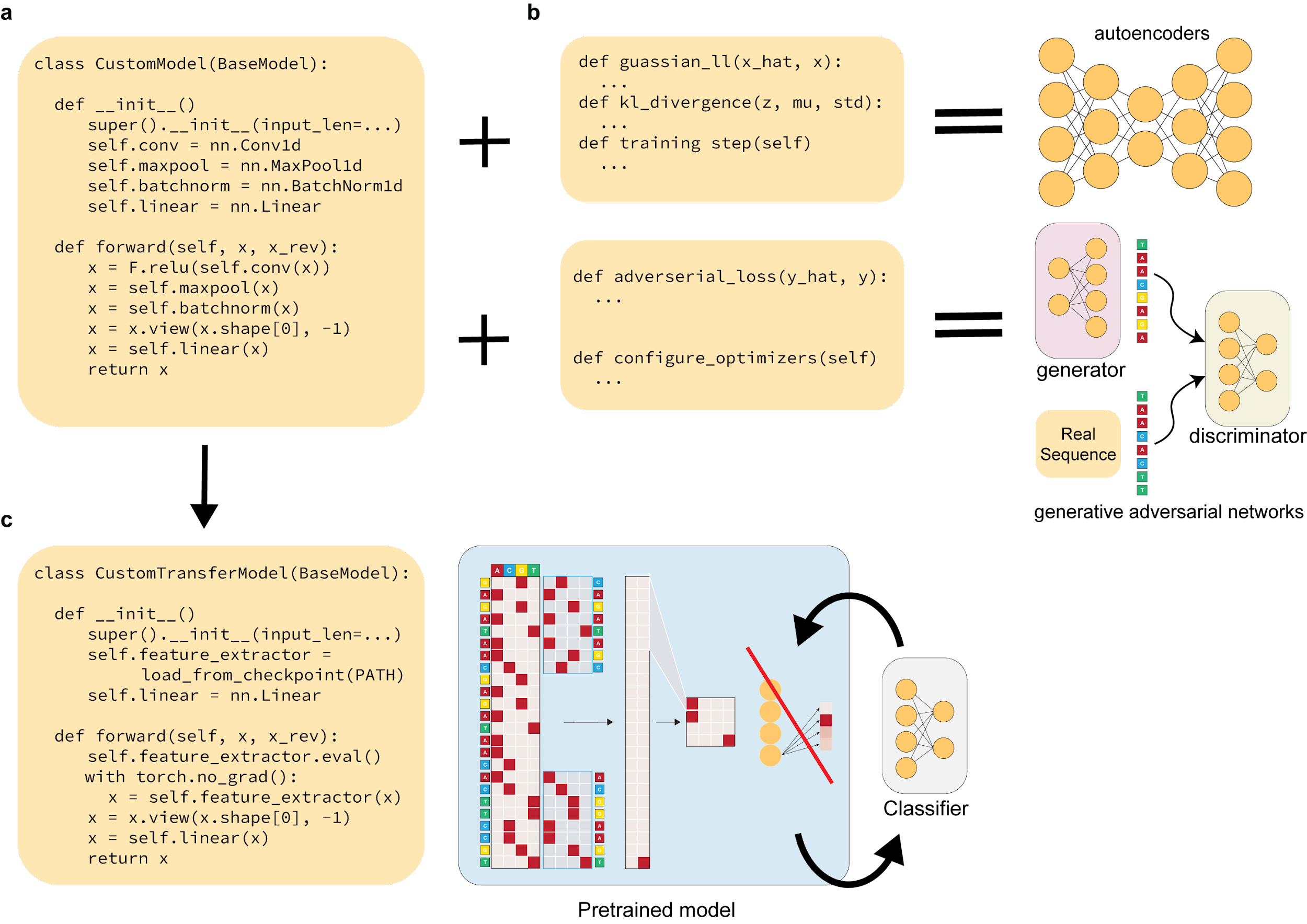

**Supplementary Figure 2. Extending EUGENe’s BaseModel to implement custom architectures. a,** Creating custom models that are compatible with EUGENe’s basic training protocol involves first inheriting from the BaseModel class (not shown), then defining the model’s architecture (__init__) and the forward propagation (forward) method. **b**, The BaseModel class can also be extended to create variational autoencoders (VAEs) or generative adversarial networks (GANs). A VAE (in its most basic form) requires creating two functions for calculating different parts of the loss and implementing how the functions are integrated into the training function. We have omitted the changes needed to define an encoder and decoder structure and how that is handled in forward. A GAN requires implementing a multipart loss function and configuring multiple optimizers to handle the training of the generator and discriminator. **c,** Transfer learning from pretrained models can be accomplished with simple changes to the initialization and forward functions. Namely, a pretrained PyTorch Lightning model needs to be loaded in the __init__ method and then utilized in the forward method.

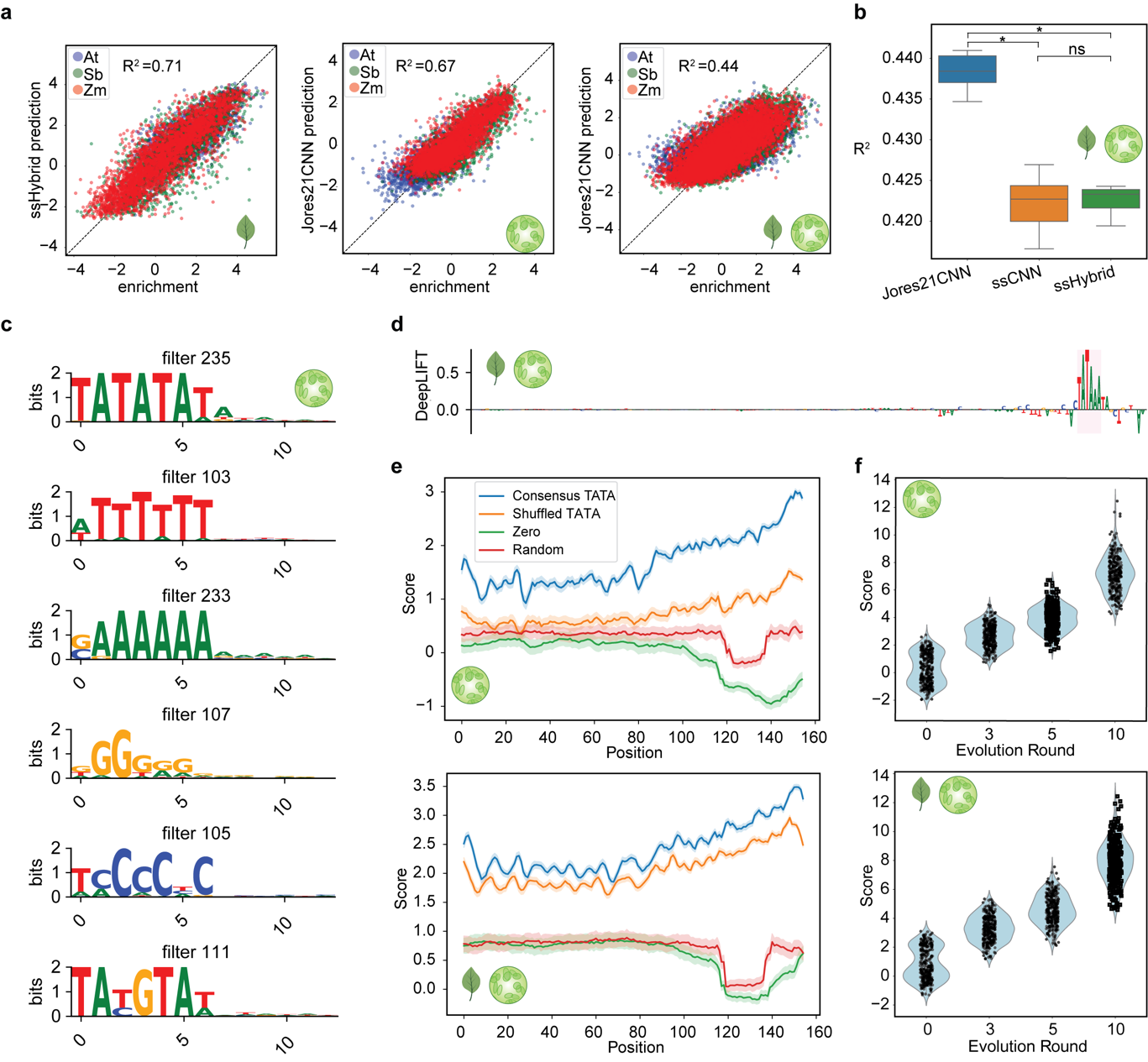

**Supplementary Figure 3. STARR-seq plant promoter activity prediction. a,** Performance scatterplots colored by species of origin for the best leaf (left), protoplast (middle) and combined (right) models. **b,** Predictive performance of all trained combined models. The boxplots show distributions of R^2^ values on held-out test data for each architecture across 5 random initializations. **c,** PWMs for a hand-selected set of learned protoplast model filters (not initialized with known PWMs). **d,** Feature attribution scores calculated using the DeepLIFT method for the sequence with the highest predicted value in the best combined model **e,** Best protoplast (top) and combined (bottom) model scores for 310 sequences with an implanted consensus TATA box motif, shuffled consensus TATA box motif, all zeros motif, and random motif at every possible position. The 95% confidence interval is shown. **f,** Model scores for the same set of 310 promoters at different rounds of evolution compared against baseline (0) for the best protoplast (top) and combined (bottom) model.

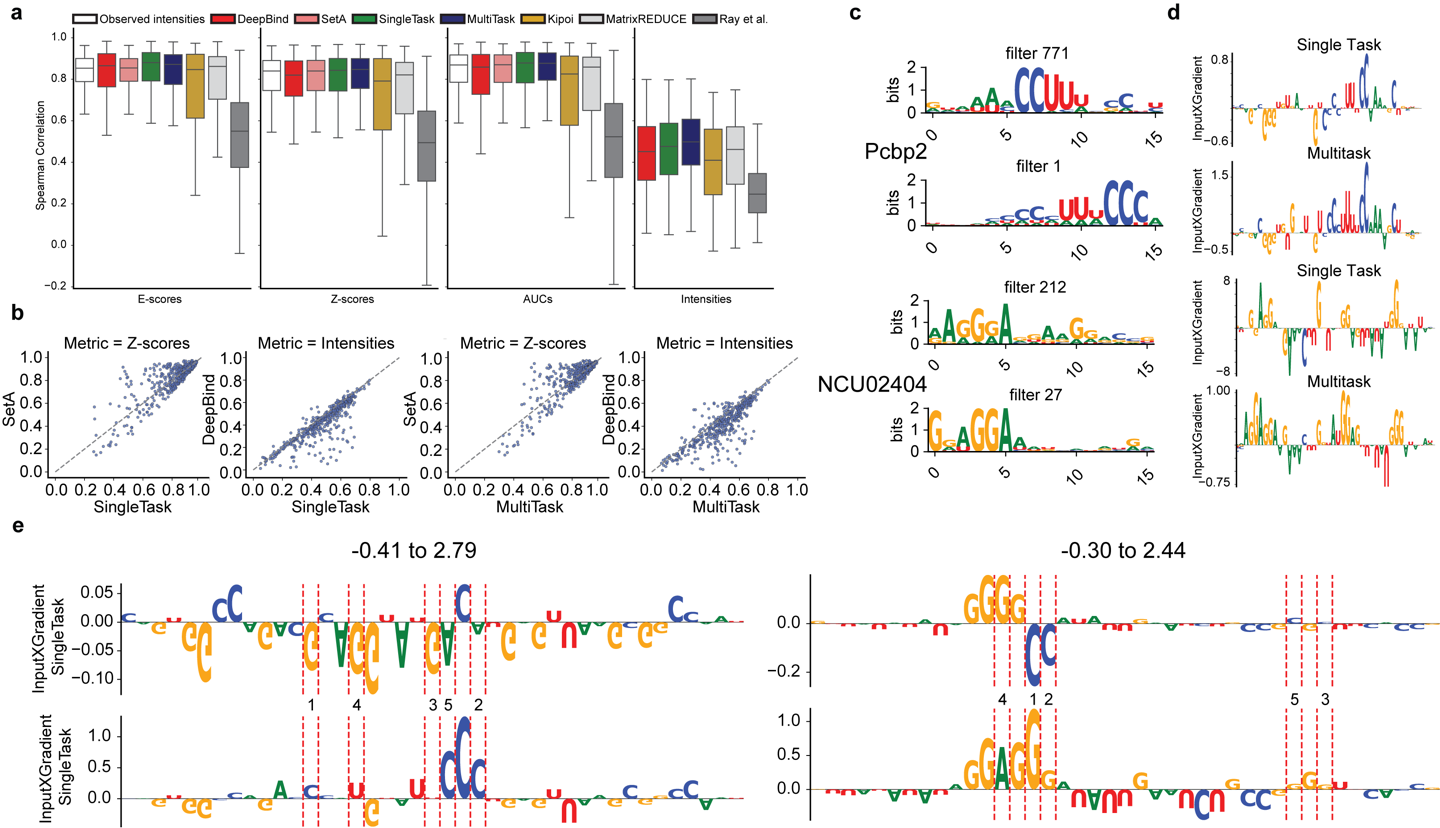

**Supplementary Figure 4. RNA binding protein (RBP) specificity prediction.** **a,** Spearman correlations across four different metrics with each metric calculated from comparisons between observed (Set B) and predicted binding intensities (see **Methods** for more details on how each metric is calculated). Each boxplot indicates a distribution of Spearman correlations across all 244 RBPs. Ray et al, MatrixREDUCE, DeepBind and Observed intensities refer to correlations calculated from predicted intensities reported in Alipanahi *et al*. Observed intensities and SetA refer to correlations calculated using the intensities from Set A probes as the predicted intensities (see **Methods**). **b,** Performance comparison scatterplots for the indicated models and metrics. Each dot indicates a comparison of the Pearson correlation between two models on a single RBP. **c,** Multitask and single task filters with TomTom significant annotations for Pcbp2 (top) and NCU02404 (bottom). **d,** The feature attributions calculated using the InputXGradient method for single task and multitask models using the sequence with the highest observed intensity in the test set for Pcbp2 (top) and NCU02404 (bottom). **e,** Two more examples of InputXGradient attribution scores for random (top row) and evolved (bottom row) sequences after evolution with the Pcbp2 (left) and NCU02404 (right) single task models. Red dashed lines indicate mutations made during evolution annotated with the round the mutation occurred in.

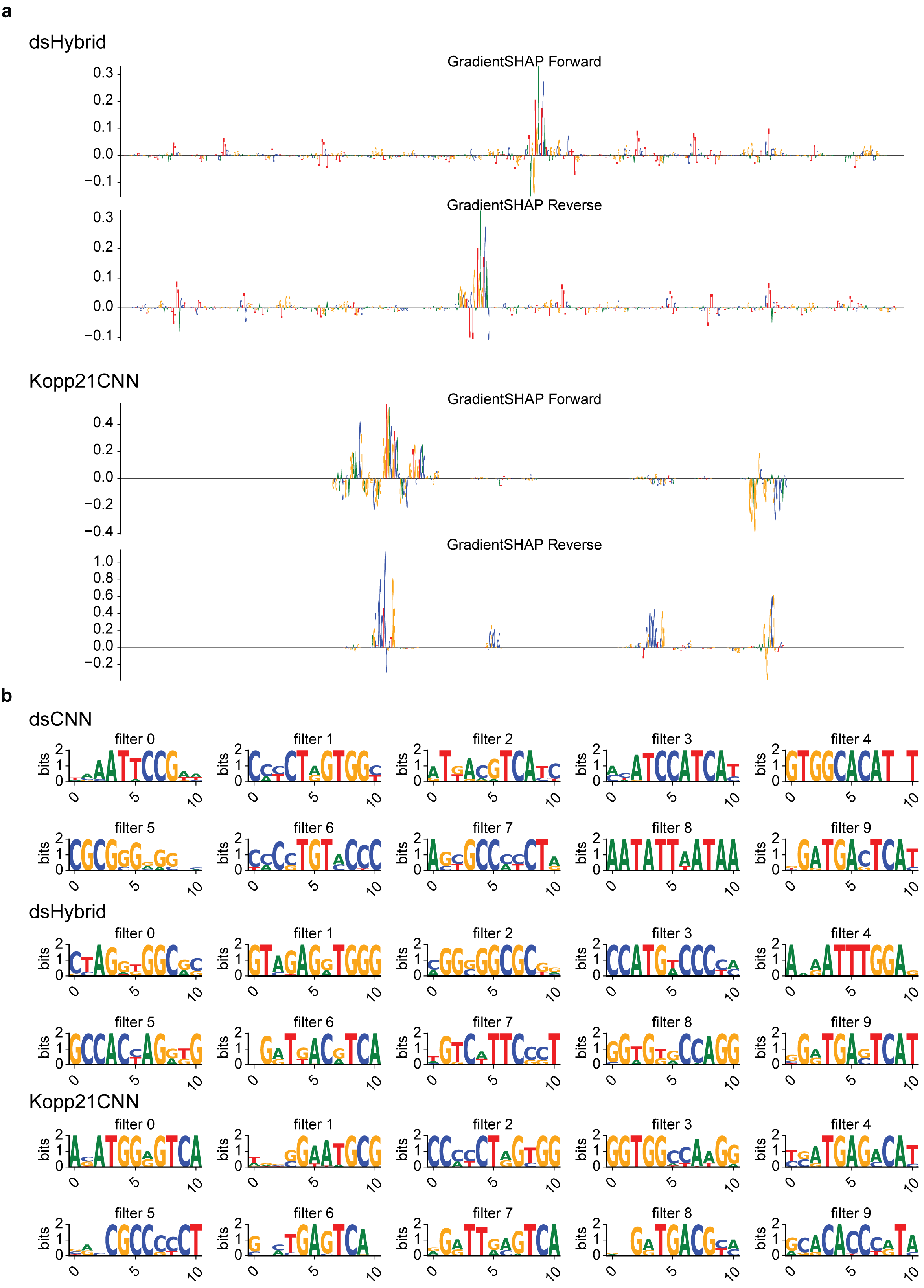

**Supplementary Figure 5. JunD binding classifier interpretation. a,** Feature attribution scores calculated using GradientSHAP for the forward and reverse complement of the sequence with the highest predictions in each of the dsHybrid and Kopp21CNN models. **b,** PWM visualizations of the 10 filters for the three convolutional architectures trained for JunD binding classification.
